# The emergence and diversification of a zoonotic pathogen from within the microbiota of intensively farmed pigs

**DOI:** 10.1101/2023.05.17.540811

**Authors:** Gemma G. R. Murray, A. S. Md. Mukarram Hossain, Eric L. Miller, Sebastian Bruchman, Andrew J. Balmer, Marta Matuszewska, Josephine Herbert, Nazreen F. Hadjirin, Robert Mugabi, Ganwu Li, Maria Laura Ferrando, Isabela Maria Fernandes de Oliveira, Thanh Nguyen, Phung L. K. Yen, Ho D. Phuc, Aung Zaw Moe, Thiri Su Wai, Marcelo Gottschalk, Virginia Aragon, Peter Valentin- Weigand, Peter M. H. Heegaard, Manouk Vrieling, Min Thein Maw, Hnin Thidar Myint, Ye Tun Win, Ngo Thi Hoa, Stephen D. Bentley, Maria J. Clavijo, Jerry M. Wells, Alexander W. Tucker, Lucy A. Weinert

## Abstract

The expansion and intensification of livestock production is predicted to promote the emergence of pathogens. As pathogens sometimes jump between species this can affect the health of humans as well as livestock. Here we investigate how livestock microbiota can act as a source of these emerging pathogens through analysis of *Streptococcus suis*, a ubiquitous component of the respiratory microbiota of pigs that is also a major cause of disease on pig farms and an important zoonotic pathogen. Combining molecular dating, phylogeography and comparative genomic analyses of a large collection of isolates, we find that several pathogenic lineages of *S. suis* emerged in the 19th and 20th centuries, during an early period of growth in pig farming. These lineages have since spread between countries and continents, mirroring trade in live pigs. They are distinguished by the presence of three genomic islands with putative roles in metabolism and cell adhesion, and an ongoing reduction in genome size, which may reflect their recent shift to a more pathogenic ecology. Reconstructions of the evolutionary histories of these islands reveal constraints on pathogen emergence that could inform control strategies, with pathogenic lineages consistently emerging from one subpopulation of *S. suis* and acquiring genes through horizontal transfer from other pathogenic lineages. These results shed light on the capacity of the microbiota to rapidly evolve to exploit changes in their host population and suggest that the impact of changes in farming on the pathogenicity and zoonotic potential of *S. suis* is yet to be fully realised.

## Introduction

Global livestock populations have grown rapidly over the past few centuries, with the global biomass of livestock now exceeding that of humans and wild mammals combined (1, 2). This has been facilitated by intensive farming systems that have also led to increased livestock population density, lower genetic diversity, and the long-distance movement of live animals. These changes are predicted to promote the emergence of pathogens (3, 4). While pathogen emergence typically arises through a pathogen jumping into a new host, pathogens can also emerge from within the microbiota already associated with a host population (5, 6). This route to pathogen emergence may be particularly important in intensive farming systems, where large population size and high population density may select for traits associated with pathogenicity while biosecurity reduces the risk of novel pathogens entering the population (7, 8).

*Streptococcus suis* was first reported as a cause of disease in farmed pigs in 1954 (9), and is now a major cause of bacterial disease in piglets and an emerging human zoonotic pathogen (10, 11). As well as being an important pathogen, *S. suis* is a ubiquitous component of the microbiota of the upper respiratory tract of all pigs. It is one of the most common bacterial species on the surface of the palatine tonsil, which is considered its main niche (12). *S. suis* disease in pigs takes the form of septicaemia with sudden death, meningitis, arthritis, and endocarditis, and most often affects piglets. It is also associated with respiratory disease, although these infections tend to be polymicrobial (13). Humans can be infected by *S. suis* either through contact with pigs or consumption of raw pork or other pig products. These infections result in similar pathologies to pigs and have high fatality rates. The first reported human case was in 1968 (14), and since then *S. suis* has led to large outbreaks in China, and has become a major cause of adult meningitis and septicaemia in South-East Asia (15–18).

While *S. suis* is a diverse species, only a small number of strains, typically characterised by multilocus sequence type or serotype, are responsible for most cases of disease (19). What determines the pathogenicity of these strains remains poorly understood despite the identification of more than 100 putative virulence genes or factors (20). Difficulties in identifying the determinants of pathogenicity in *S. suis* have been attributed to its complex pathogenesis and high level of genetic diversity (20). Few studies have considered virulence factors in strains other than sequence type (ST) 1, which is responsible for most cases of *S. suis* disease in both pigs and humans worldwide (19).

In this study, we carried out a population-genomic analysis of 3,070 bacterial isolates sampled from tonsil and nasal swabs of pigs and wild boar, and blood and sites of infection in pigs and humans with *S. suis* disease, from Europe, North America, Asia and Australia, dating from 1960 to 2020, to investigate the emergence, diversification and geographic spread of pathogenic lineages of *S. suis*. Through development of a new whole-genome typing schema we identified 10 pathogenic lineages with broad geographic distributions, dated their origins, and mapped their movements between countries. We identified genomic changes associated with the emergence of these lineages and investigated their origins. We also considered the impact of pathogenicity on broader evolutionary dynamics, the ongoing diversification of pathogenic lineages, and how farming practices may have contributed to these processes.

## Results

### Pathogenic lineages emerged from a subpopulation of a diverse and largely commensal species

We sequenced the genomes of isolates identified in laboratory assays as *Streptococcus suis* from pigs from farms in Denmark (n=173), Germany (n=166), the Netherlands (n=168), Spain (n=200), the UK (n=49), Myanmar (n=701) and the USA (n=293) (Table S1). These isolates date from 1960 to 2020. Those from Denmark, Germany, the Netherlands and Spain included similar numbers of isolates from the tonsils or noses of pigs without *S. suis*-associated disease (carriage isolates; n=188), and isolates from blood or sites of infection in pigs with respiratory (n=196) or systemic (n=196) forms of *S. suis*-associated disease (disease isolates). Those from Myanmar only included carriage and environmental isolates and those from the USA only disease isolates. The collection from Spain included isolates from a wild boar population (n=34). We combined these with previously published genome sequence data from isolates from Australia (n=143), Canada (n=200), China (n=217), Denmark (n=1), Vietnam (n=191), the UK (n=441), the Netherlands (n=101), Spain (n=10) and the USA (n=16). This led to a collection of 3,070 high-quality genome assemblies, including 48 complete genomes.

To investigate the population structure of *S. suis* we considered genetic variation estimated from both single nucleotide polymorphisms (SNPs) in genes that are present across all isolates (core genes) and from the presence/absence of accessory genes (Figure 1a, S1-4). Both revealed high levels of diversity and distinguished a cluster of closely related isolates that includes most of our collection (2,424/3,070 isolates). Isolates in this cluster have a maximum pairwise distance of 0.08 differences per nucleotide in a core genome alignment, while the remaining 647 isolates diverge from members of this cluster by between 0.13 and 0.32 differences per nucleotide site. We refer to this cluster as the ‘central population’ of *S. suis* (a previous study referred to it as ‘normal’ *S. suis* (21)). The isolates that fall outside of the central population form several distinct clades in a core genome phylogeny (Figure S1). While it has been suggested that these may represent distinct species, a previous study concluded that phenotypic similarities and extensive gene exchange between some of these lineages and the central *S. suis* population meant that there was insufficient evidence to assign them to a new species or subspecies (21). In support of this conclusion, we found that divergent isolates are less clearly distinguished from the central population of *S. suis* by differences in gene content than by SNPs in core genes (Figure 1a, S1).

**Figure 1.**
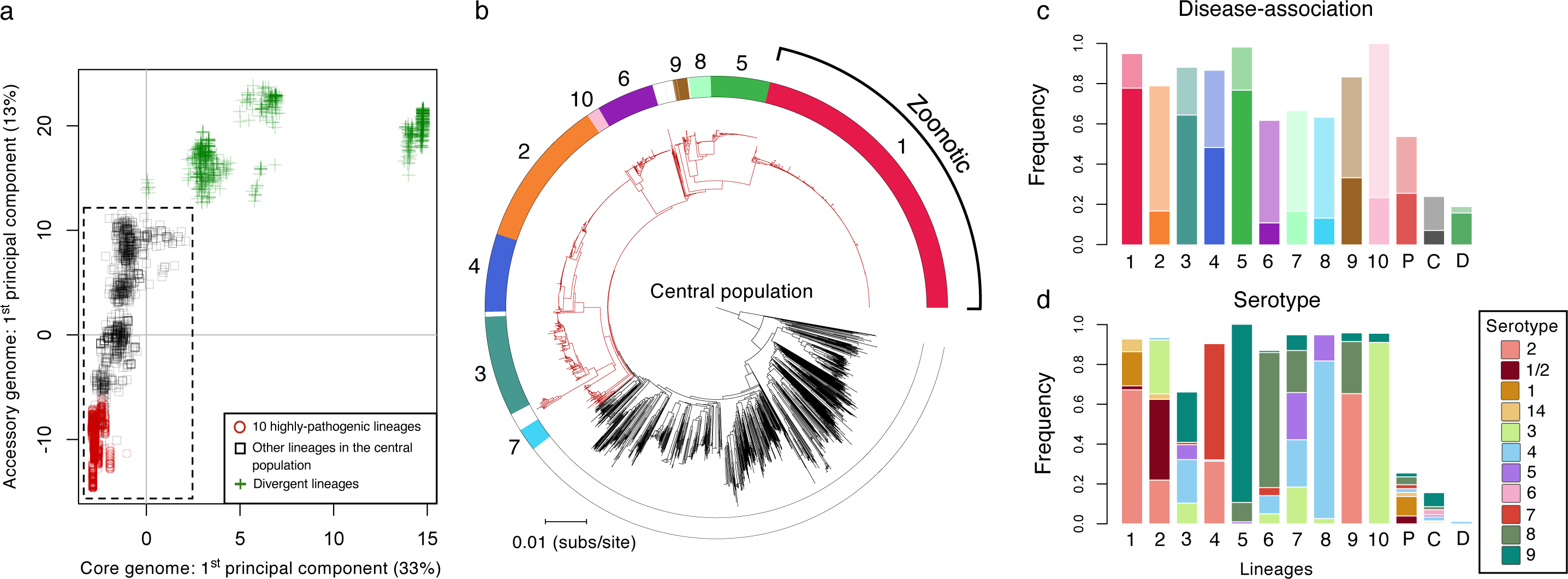
The relationship between pathogenicity and genetic structure within *Streptococcus suis*. (a) The first components of principal component analyses of the presence/absence of accessory genes and SNPs in core genes plotted against one another for all 3,070 isolates in our collection (other components and their eigenvalues are shown in Figures S3 and S4). Points represent individual isolates and different shapes/colours represent categories of lineages. Isolates in the dashed box fall within the central population of *S. suis*. (b) A core genome phylogeny of the central population of *S. suis* (excluding divergent isolates). Branches in the pathogenic clade are coloured red and branches outside of this clade are coloured black. Colours in the outer ring indicate the 10 most common lineages in our collection, which we also find to be the most pathogenic. Lineage 1 largely corresponds to ST1, which is associated with most cases of zoonotic disease in humans. (c) The frequency of disease-associated isolates in each of the 10 pathogenic lineages (1–10), in other lineages within the pathogenic clade (P, coloured red in (b)), in the central population outside of the pathogenic clade (C, coloured black in (b)), and in divergent lineages (D, green addition signs in (a)). Bars 1-10 are coloured to match the outer ring in (b) with the lower part of the bar (deeper colour) representing the frequency of systemic disease isolates and the upper part of the bar (paler colour) representing the frequency of respiratory disease isolates. (d) The frequency of disease-associated serotypes in the same groups as shown in (c).

To mitigate any geographic bias, we characterised variation in disease-association across *S. suis* using a subset of our collection that includes isolates that (a) have a well-characterised association with disease or carriage states, and (b) are from a country from which we have large samples of both disease and carriage isolates in our collection (>40 of each). This subset includes isolates from Denmark, Germany, the Netherlands, Spain, Canada, and the UK (n=1,193). In agreement with previous studies, our analysis revealed that while disease isolates are present across the entire genetic diversity of *S. suis* (including in divergent clades) they are concentrated in a subpopulation of the central population (Figure 1b,c) (22).

Using variation in both core and accessory genes we partitioned the central population of *S. suis* into clusters of closely related isolates (lineages) (Table S1). A reference database describing these lineages, which will allow for classification of further isolates using the same schema, is available online (www.bacpop.org/poppunk). Combining this with data on disease-association, allowed us to identify 10 ‘pathogenic’ lineages. In each of these lineages at least 60% of the isolates from the subset of our collection described above are disease-associated, compared to 26% of isolates from other lineages in the central population, and 19% of isolates from divergent lineages (Figure 1c). Combined, the 10 lineages account for 80% of disease-associated isolates in our collection (Figure 1c, Table 1) (19). These lineages also have much higher frequencies of serotypes that are commonly associated with disease: 88% of isolates from these 10 lineages have a disease-associated serotype (1, 1/2, 2, 3, 4, 5, 6, 7, 8, 9 or 14) compared to 16% from other lineages (Table 1, Figure 1d).

**Table 1.**
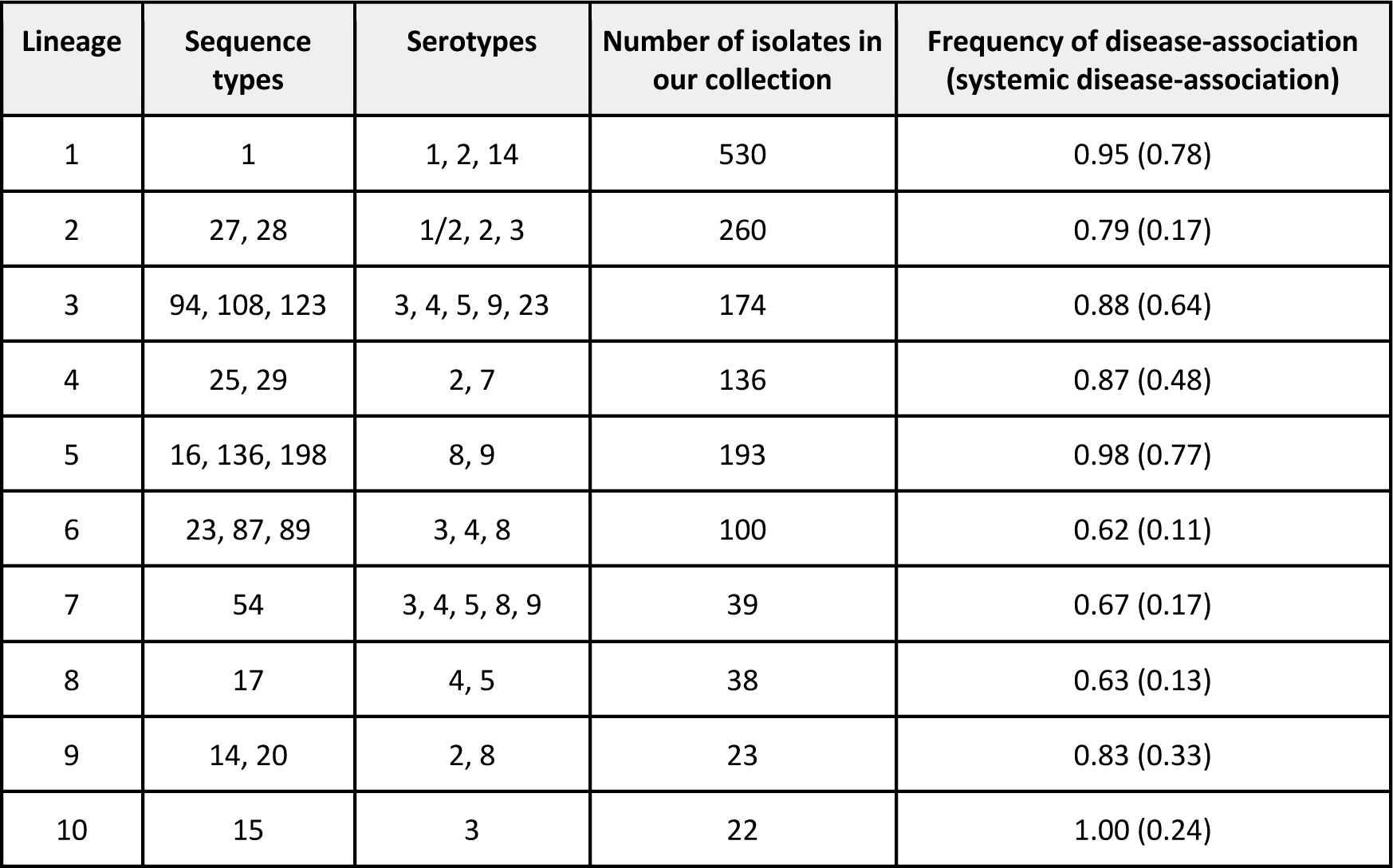
Sequence types and serotypes associated with the 10 pathogenic lineages. Each of the 10 pathogenic lineages include multiple sequence types and serotypes. Sequence types and serotypes present in >5% of the isolates from our collection of each lineage are shown here, along with the number of isolates from these lineages in our collection, and the frequency of disease-association (and systemic disease-association) in the subset of our collection we used to characterise pathogenicity. A description of the sequence types and serotypes of all isolates in our collection is provided in Table S1.

The 10 pathogenic lineages of *S. suis* fall within a subpopulation of the central *S. suis* population. Core nucleotide distances between these 10 lineages are, on average, lower than between lineages across the rest of the central population (Figure S1, S2). Nevertheless, they are not clearly distinguished from other lineages in the central population; there are an additional 34 lineages (98 isolates) that fall within the clade that includes the 10 pathogenic lineages (Figure 1b). Combined, these 34 lineages show an elevated rate of disease-association: 54% of isolates are disease-associated compared to 26% from the central *S. suis* population outside of this clade (Figure 1c). They are also enriched for pathogenic serotypes; 41% of isolates have a pathogenic serotype compared to 8% in other lineages of the central population (Figure 1d). This suggests that some of these lineages may also be pathogenic, but we cannot confidently characterise them as such due to small sample sizes (all have <15 isolates in the subset of our collection used for characterising disease-association).

### Consistent genomic changes reveal evolutionary constraints on pathogen emergence

Previous studies that have investigated genes associated with pathogenicity in *S. suis* have tended to focus on isolates from the zoonotic lineage ST1, which is both highly pathogenic in pigs and responsible for most cases of human disease. Here, we aimed to identify genes associated with pathogenicity more broadly across *S. suis* (i.e. across all pathogenic lineages of *S. suis*). To do this, we identified genes that are present at >70% higher frequency in isolates from any of the 10 pathogenic lineages relative to both lineages from the central *S. suis* population outside of the pathogenic clade and divergent lineages. While no genes are uniquely present in pathogenic lineages, a small number (n=37) are present at >70% higher frequencies in isolates from pathogenic lineages than both isolates from other lineages from the central population and from divergent lineages (Figure S5, Table S2). Most of these genes (26/37) fall within just three genomic islands: Island 1 (SSU_RS05400-SSU_RS05325 in the published annotation of P1/7), Island 2 (SSU_RS02325-SSU_RS02355) and Island 3 (SSU_RS01130-SSU_RS01185) (Figure 2, Figures S6-8, Table S2). Five of the 11 genes outside of the three genomic islands have previously been described as associated with pathogenicity in *S. suis*, these are *mrp*, *sspA*, *sspep*, SSU0587 and *sstgase* (20).

**Figure 2.**
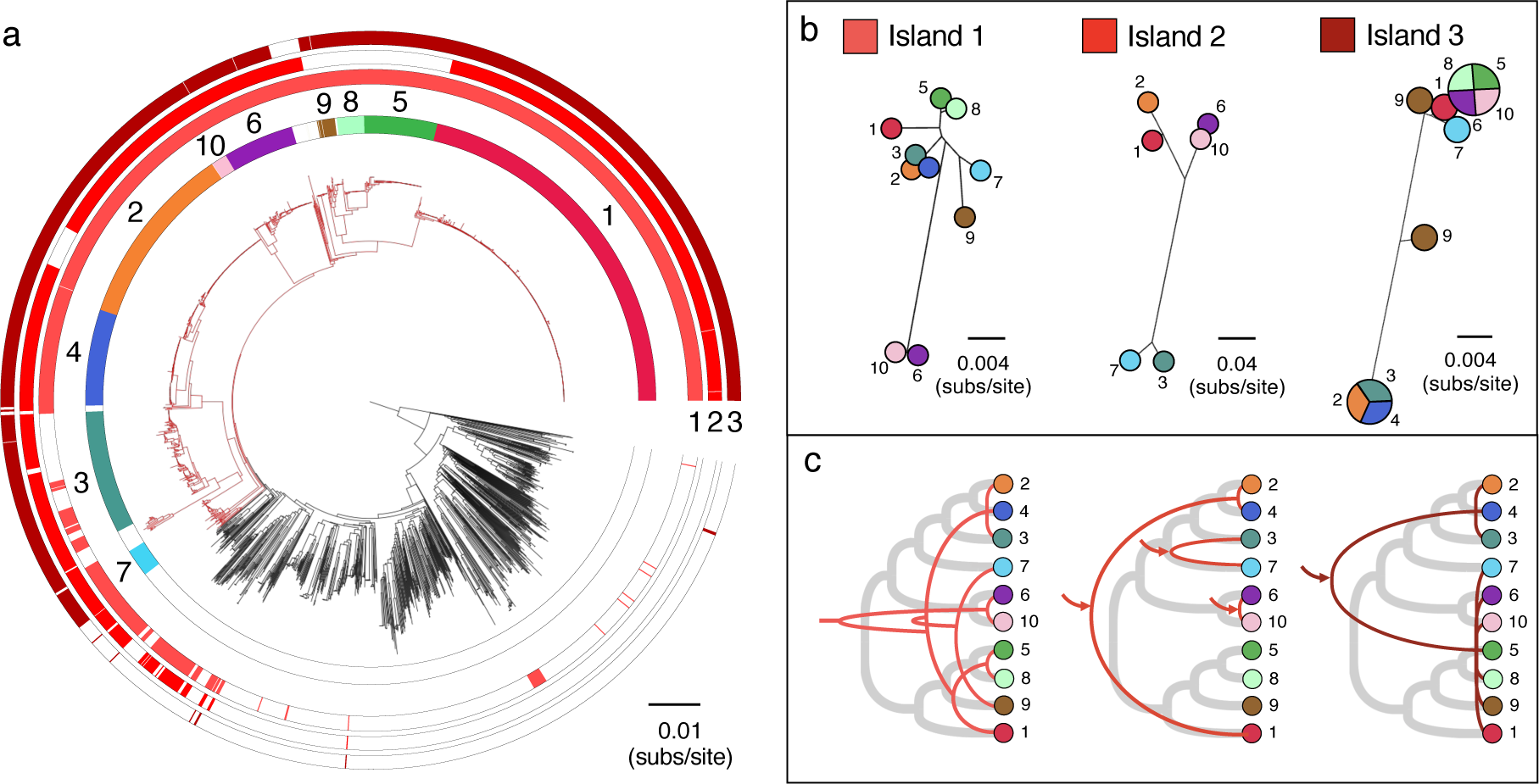
Divergent evolutionary histories of the three pathogenicity-associated genomic islands. (a) The same core genome phylogeny as shown in (Figure 1b) with three additional outer rings describing the presence of the three pathogenicity-associated genomic islands (1, 2 and 3). (b) Trees representing median genetic distances between pathogenic lineages based on coding regions of each of the three genomic islands (as described in Figures S6-S8). For Island 3, lineages that have average distances of zero are represented by the same circle split into segments and lineage 9 is represented by two circles due to the presence of two divergent versions of the island. (c) Cladograms representing inferred patterns of acquisition and inheritance of these three islands (red lines) compared to the core genome (thicker grey lines). Arrows indicate inferred acquisitions from outside of *S. suis*. A single inferred recombination event is omitted from the description of the history of Island 3 shown in (c).

Island 1 is present in >95% of isolates from 9/10 pathogenic lineages, only 17% of isolates from the central population outside of the pathogenic clade (and these tend to be from lineages closely related to the pathogenic clade; Figure 2a), and <1% of isolates from divergent lineages. The island contains genes predicted to encode proteins involved in the breakdown of host glycans present in the extracellular matrices of mammalian tissues. It includes a putative heparan sulphatase (SSU_RS05330 in P1/7) and a putative hyluronate lyase (SSU_RS05335-SSU_RS05350 in P1/7) which cleave hyaluronic acid found in all connective body tissue. However, in zoonotic lineage 1 the gene encoding the hyluronate lyase protein is always truncated, likely leading to loss of function. The phosphotransferase system and other enzymes present in this island may be associated with sugar transport and metabolism of host aminoglycans which are degraded extracellularly by *S. suis*. The degradation of extracellular matrix proteins may both contribute to the spread of *S. suis* through the connective tissue and provide a source of energy at epithelial surfaces.

Nucleotide diversity within Island 1 is similar to and broadly correlated with diversity in core genes both within and between pathogenic lineages (Figures 2, S9, S10). The island is also found next to the same core gene (*nrdF* or SSU_RS05320 in P1/7) in 97% of isolates. Together this suggests that the island was acquired once by a common ancestor of all pathogenic lineages. As homologous recombination is common in *S. suis* (23), the inheritance of this island is unlikely to have been entirely vertical, but our observations suggest that it has followed similar patterns to core genes (Figure 2c).

Island 2 is present in 84% of isolates from the 10 pathogenic lineages, and only 11% from the central population outside of the pathogenic clade, where it tends to be carried by isolates that are both closely related to the pathogenic clade and also carry Island 1 (Figure 2a). It is not found in any isolates from divergent lineages. Its presence is variable across pathogenic lineages: it is present in >80% of isolates from lineages 1, 2, 3, 4, 6, 7 and 10, but absent from lineages 5, 8 and 9. Island 2 encodes a major pilin subunit (SSU_RS02345), a minor pilin subunit (SSU_ RS02335 and SSU_RS02340), and a pilin specific sortase (SSU_RS02350) that is involved in maturation. The minor pilin subunit (SSU_ RS02335 and SSU_RS02340) was previously described as a pseudogene in several pathogenic strains of *S. suis* (including P1/7) (24). We find evidence that this is true of all isolates in lineage 1 and 2, and several from other lineages due to premature stop codons. Nevertheless, Fittipaldi *et al.* (24) showed that despite this truncation, *S. suis* P1/7 still produces a major pilin subunit. While they found that this did not play a role in adherence to porcine brain microvascular endothelial cells or virulence in a mouse model of sepsis, Faulds-Pain et al. (25) later showed that the pilin specific sortase in this island was essential for causing disease in pigs via the intranasal route of infection. We find that in a small number of isolates (largely from lineages 3 and 7) the major pilin subunit is truncated (see Figure S6 for an example). Further work is required to understand the impact of this on the function of the island.

Diversity within Island 2 suggests multiple acquisitions from outside of *S. suis*: three divergent versions are carried by pathogenic lineages (lineages 1 and 2 carry a similar version that is distinct from a version carried by lineages 6 and 10, and a version carried by lineages 3 and 7; Figure 2b,c, Figure S9). The island is found next to the same core gene (*murD* or SSU_RS02355 in P1/7) in 99% of isolates and divergence within each of the pathogenic lineages is similar to and correlated with diversity in both core genes and Island 1 (Figure S9, S11). Divergence between the pairs of lineages that carry each of the three versions of the island is variable, and similar to distances between these lineages based on core genes and Island 1. This suggest that the presence of each version of Island 2 in a pair of pathogenic lineages is a consequence of a single acquisition by a common ancestor. As the most recent common ancestor of lineages 1 and 2 is also that of the entire pathogenic clade, this suggests that the version of this island carried by lineages 1 and 2 was maintained since the common ancestor of the pathogenic clade, while the version carried by lineages 6 and 10 has been acquired much more recently.

Island 3 shows the strongest association with pathogenic lineages. It is present in >95% of isolates in the 10 of the pathogenic lineages and only 1% of isolates from the central population outside of the pathogenic clade. It is entirely absent from divergent lineages. It contains genes that code for an ABC transporter, a ROK (repressor, open reading frame, kinase) family gene (26), and a large predicted surface protein with similarity to a PTSII subunit. The ROK is similar to an *E. coli* repressor (Mlc), which represses several genes including two phosphotransferase system (PTS) genes (27). In *Salmonella Typhimurium* Mlc positively regulates expression of the *Salmonella Typhimurium* pathogenicity island 1 genes by reducing the expression of the negative regulator HilE (28).

There are two divergent versions of Island 3 in our collection: one carried by lineages 1, 5, 6, 7, 8, 9 and 10, and the other by lineages 2, 3 and 4. While some isolates from lineage 9 carry a version that differs from these two, this appears to be the result of recombination between the two versions. In these isolates there are extended tracts of SNPs that are shared with either of the two main versions and there is only one unique SNP. Island 3 is found next to the same core gene in 98% of isolates in our collection (SSU_RS01115 in P1/7) and diversity in the island is positively correlated with diversity in core genes within lineages (Figure S12). Within-lineage diversity is generally lower in this island than in the other two islands (Figure S9), but this may reflect stronger selective constraint rather than a more recent origin. This is supported by lower divergence at 1st and 2nd codon positions relative to 3rd codon positions (Figure S13). Together this suggests that the presence of Island 3 in each of the pathogenic lineages tends to be the result of a single acquisition by a common ancestor of the lineage. We find evidence of only one case of a second acquisition event by a pathogenic lineage: one isolate from lineage 2 carries the version of Island 3 associated with lineages 1, 5, 6, 7, 8, 9 and 10 (PH2016-139). While very low diversity within the two versions of the island means that the exact relationships between those carried by different lineages cannot be reconstructed, it indicates that for most pathogenic lineages Island 3 was obtained from another pathogenic lineage, and that the transfer did not occur long before the most recent common ancestor of each lineage.

Each of the three pathogenicity-associated islands are sometimes present in lineages other than the 10 pathogenic lineages (Figure 2a, Table S1). All three are present together in only six additional lineages (15 isolates): lineages 18, 107, 144, 164, 466 and 511. Of these, three fall inside the pathogenic clade and three within the central population but outside of the pathogenic clade. While the three in the pathogenic clade are all represented by multiple isolates, the three outside the pathogenic clade are all only represented by a single isolate. Combined, the six lineages have a broad geographic distribution that spans Europe, North America, Australia and Asia. Lineage 18 alone includes isolates from Canada, the USA and Myanmar, that date from 1987 to 2019. Some of these lineages may represent additional pathogenic lineages that are not well represented by our collection, while others may represent the chance acquisition of these islands by isolates that are otherwise ill-suited to a pathogenic ecology.

### The emergence and spread of pathogenic lineages are linked to growth in pig farming and international trade

We used the temporal structure in our collection of isolates from the six most common pathogenic lineages to construct dated phylogenies (Figures 3, S14). These reveal that the dates of the most recent common ancestors of these lineages range from 1827 (lineage 2; 95% HPD: 1798-1854) to 1951 (lineage 5; 95% HPD: 1944-1958) (Table S3). These origins largely predate a rapid period of growth in pig numbers in several European countries that accompanied the wide-scale shift to larger farms and indoor rearing in the early 20^th^ century (29, 30), and instead accompany a period of human population growth in Europe and North America that followed the Industrial Revolution (31) (Figure S15).

**Figure 3.**
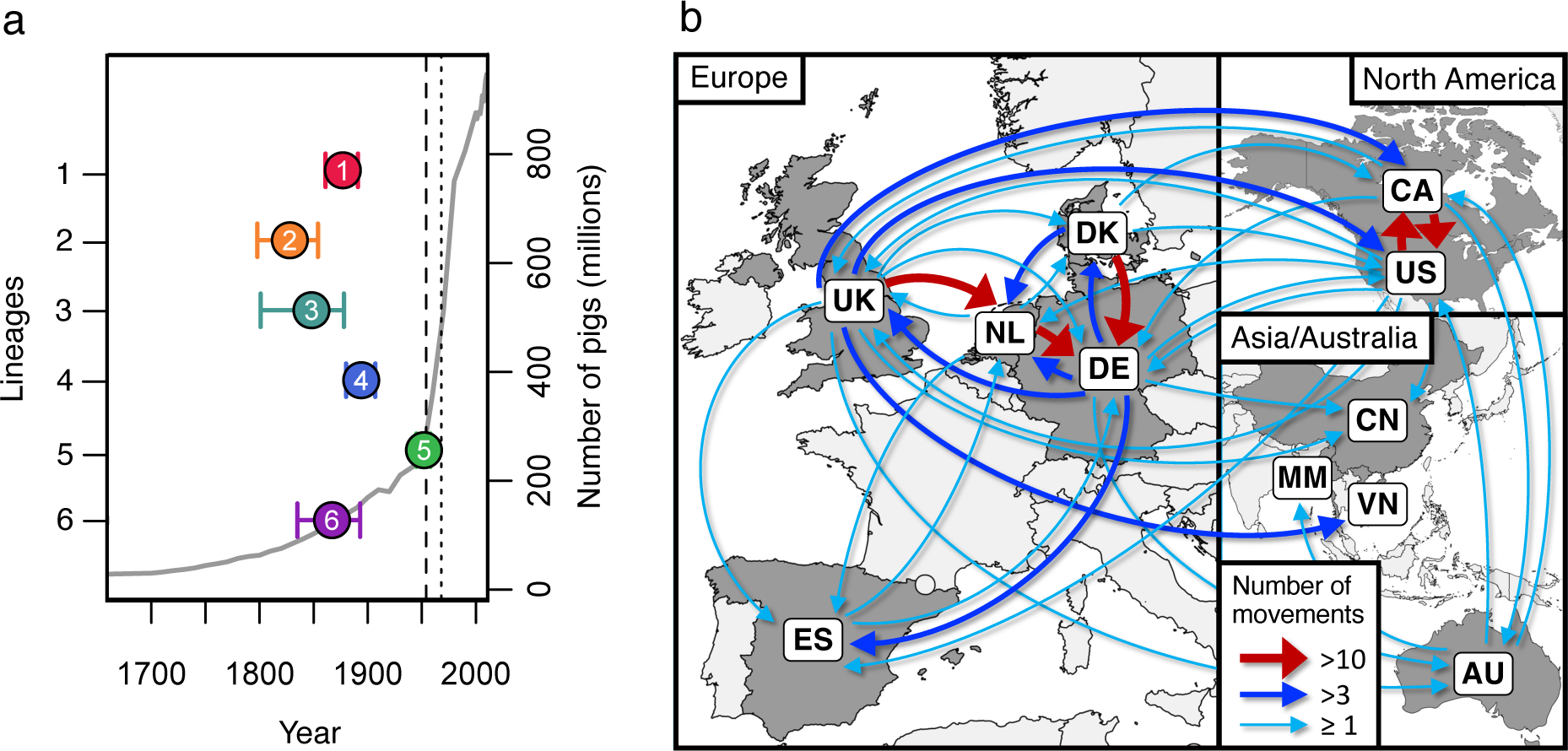
Dates of emergence and paths of between-country transmission for the six most common pathogenic lineages. (a) Estimates of the dates of the most recent common ancestors of the six most common pathogenic lineages (coloured points) against an estimate of the global number of pigs (grey line). Country-specific estimates of pig numbers are shown in Figure S15. The vertical dashed line shows the date of the first reported case of *S. suis* disease in pigs (1954) and the dotted line shows the first reported human case (1968). (b) Map showing inferred routes of transmission of these six pathogenic lineages between the countries in our collection. Arrows represent routes with at least one inferred transmission event. Routes with more than ten inferred transmission events are shown in red, those with more than three in blue, and those with one to three in turquoise. Further details of the numbers and rates of movements between countries across our six lineages are shown in Figures S18-20 and Table S4.

The 10 pathogenic lineages all have a broad geographic spread: each are found in at least 6/11 countries in our collection. Similarly, outside of these lineages we find little evidence of geographic structure in *S. suis*: isolates from each well-sampled country span the diversity of *S. suis* (Figure S16). This is also true of isolates in our collection from a wild boar population in Spain (Figure S17). Using our dated phylogenies and a discrete asymmetric model we estimated the number of between-country movements for each of the six most common pathogenic lineages and their relative rates. We estimated between 16 and 45 between-country movements for each lineage, and 182 across all six lineages (Table S4). This is likely to be a substantial underestimate of the true number of movements due to incomplete sampling.

Across all lineages, our highest inferred rate of between-country transmission was from Denmark to Germany, followed closely by Canada to the USA, the Netherlands to Germany, and the USA to Canada (Figures 3 and 4, S18, Table S4). This is consistent with these movements being driven by international trade in live pigs; over the last 80 years the top three global exporters of live pigs have been Denmark, the Netherlands and Canada and top two importers have been Germany and the USA (www.fao.org/faostat). Additionally, we find no evidence of the transmission of pathogenic lineages into Australia since its ban on live pig imports in the mid-1980s (32). While we infer several transmission events from Europe and Canada to Australia, the most recent is a movement of zoonotic lineage 1 from the UK that we estimate to have occurred between 1981 [95% CI: 1976-1987] and 1986 [95% CI: 1981-1991].

**Figure 4.**
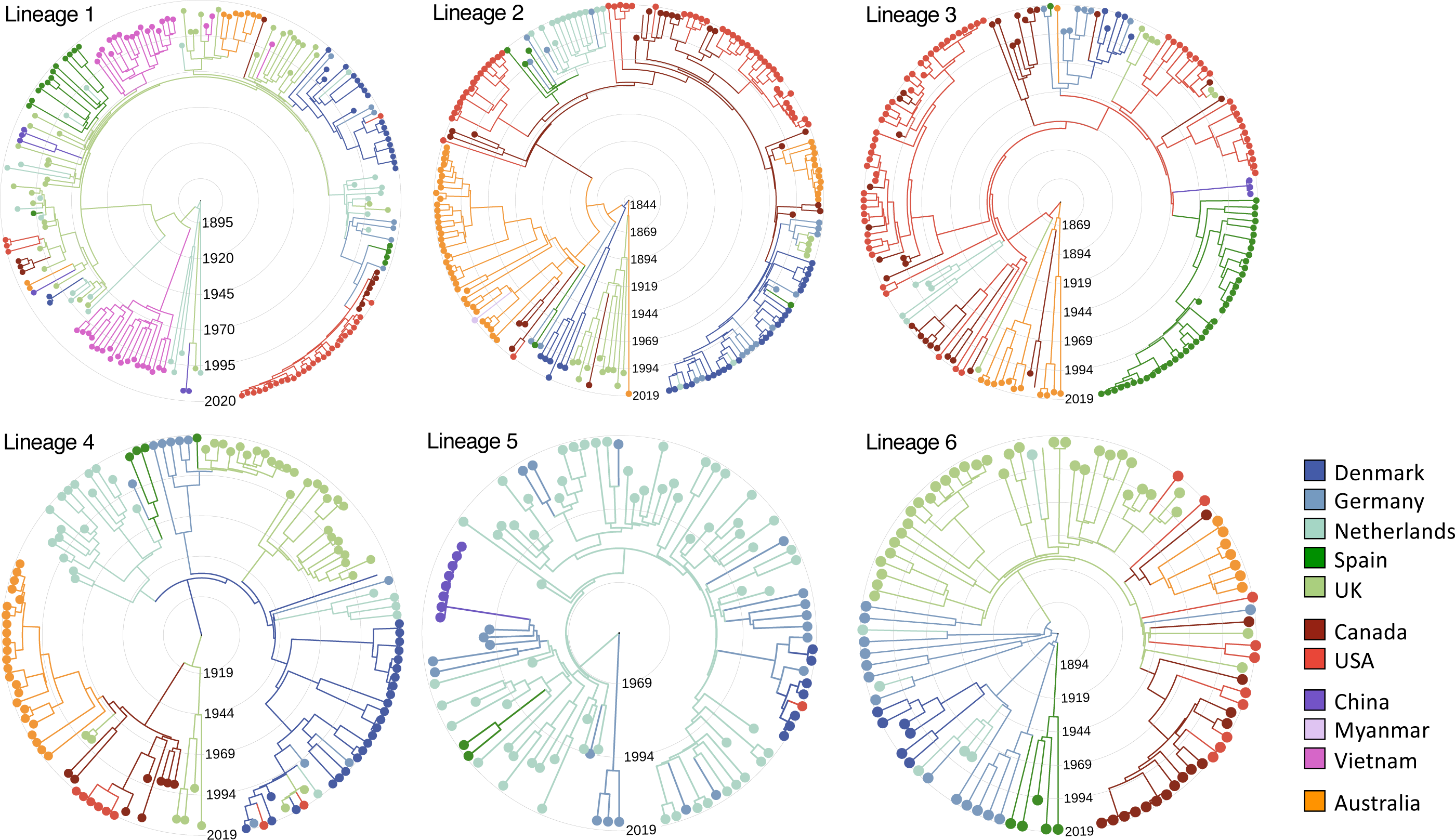
Reconstructions of the emergence and spread of the six most common pathogenic lineages. Time-scaled phylogenies for the six most common pathogenic lineages of S. suis coloured by country of origin. Colours of branches represent the most likely ancestral location inferred from ancestral state reconstructions with an asymmetric discrete transition model in BEAST.

We also observe a an unusually low frequency of pathogenic lineages in our collection from Myanmar. Pig farming in Myanmar is typically small-scale and most of our samples from this country are from backyard or semi-intensive farms that typically have 10-30 pigs. While all of our isolates from Myanmar are non-clinical and so we would expect a low frequency of pathogenic lineages, they are less likely to be from pathogenic lineages than non-clinical isolates from other countries: <1% of isolates from Myanmar compared to a mean of 18% across other countries (Table S5). In fact, none of our isolates from backyard and semi-intensive farms in Myanmar are from pathogenic lineages (0/109 isolates from the central *S. suis* population); the only pathogenic lineage isolate in our collection is from a larger commercial farm (1/39) with around 2,000 pigs.

As well as the spread of pathogenic lineages between pig farms in different countries, we find evidence of transmission between farmed pigs and wild boar. The two isolates from our collection of Spanish wild boar that are from a pathogenic lineage are both from lineage 1. They form a single clade that falls within a clade that appears to have been circulating in Spain since the 1970s. This indicates recent transmission between farmed pigs and local wild boar.

Individual lineages show different patterns of between-country transmission (Figure 4, S19, S20). This may reflect variation in both the prevalence of these lineages and their dates of introduction to particular countries. In particular, we find evidence that zoonotic lineage 1 has been circulating in the UK and the Netherlands for at least 50 years, with frequent transmission between these two countries, while the earliest evidence we find of its presence in North America is 2009 [95% CI: 2007-2011]. This is consistent with this lineage both being less commonly associated with disease in North America, and reports of an increase in its rate of detection over the last few years. In contrast, lineage 2 shows evidence of circulation in Canada for around 80 years and repeated transmission from Canada to the USA. Unfortunately, uncertainty in the inferred locations of the common ancestors of each of the lineages means that we cannot confidently infer the country of origin for any pathogenic lineage.

### Pathogenic lineages have variable associations with systemic disease and carry unique genes

Genetic and phenotypic diversity can make it more difficult to control the spread of a pathogen. For example, it can lead to difficulties in developing cross-protective vaccines. It can also make a pathogen more capable of evolving to evade control measures. In *S. suis* we find evidence of both phenotypic and genetic variation between pathogenic lineages. The six most common pathogenic lineages in our collection vary in their frequency of both association with disease relative to carriage (Chi-Squared test, *p* = 3.6 x 10^-8^) and systemic disease relative to respiratory disease (Chi-Squared test, *p* = 2.0 x 10^-20^). Pairwise comparisons reveal two distinct groups: lineages 1, 3, 4 and 5 have higher frequencies of association with systemic disease while lineages 2 and 6 have higher frequencies of association with respiratory disease (Figure S21). This is consistent with previous studies that found that ST28, which is part of lineage 2, is less virulent in mouse models than both ST1 (lineage 1) and ST25 (lineage 4) (33).

Pathogenic lineages also show evidence of variation in their evolutionary dynamics and genome content. Average rates of nucleotide substitutions per site were highest for lineage 5, while the ratio of transitions to transversions is lower for lineages 2 and 4 (Figure S22). The six most common pathogenic lineages can be distinguished by their carriage of 2-32 lineage-specific genes (that are present in >95% of isolates in that lineage and <5% of isolates from other lineages; Table S6). For instance, 17 genes distinguish zoonotic lineage 1. Far more genes are associated with multiple pathogenic lineages; 309 genes are present in >95% of isolates in at least one of the six most common pathogenic lineages and in <5% of isolates outside of the pathogenic clade (Table S6). Apart from lineages 2 and 4 that share more genes than other pathogenic lineages, there is little evidence of clustering of lineages by shared gene content (Figure S23).

### Pathogenic lineages continue to diverge from commensal lineages and diversify

Pathogenic lineages of *S. suis* tend to have smaller genomes than those from outside of the pathogenic clade, and pathogenic lineages with higher frequencies of association with disease tend to have smaller genomes than those with higher frequencies of association with carriage (Figure 5a). This pattern was previously described in a study based on a smaller sample of *S. suis* isolates (22). Using the temporal structure in our collection, we additionally found that older isolates of pathogenic lineages tend to have larger genomes than more recent isolates. This suggests a gradual and ongoing process of genome reduction in pathogenic lineages (Figure 5b). While all pathogenic lineages show this pattern, we observe variation in the rate of genome reduction across lineages, with zoonotic lineage 1 showing the slowest rate and lineage 5 the fastest.

**Figure 5.**
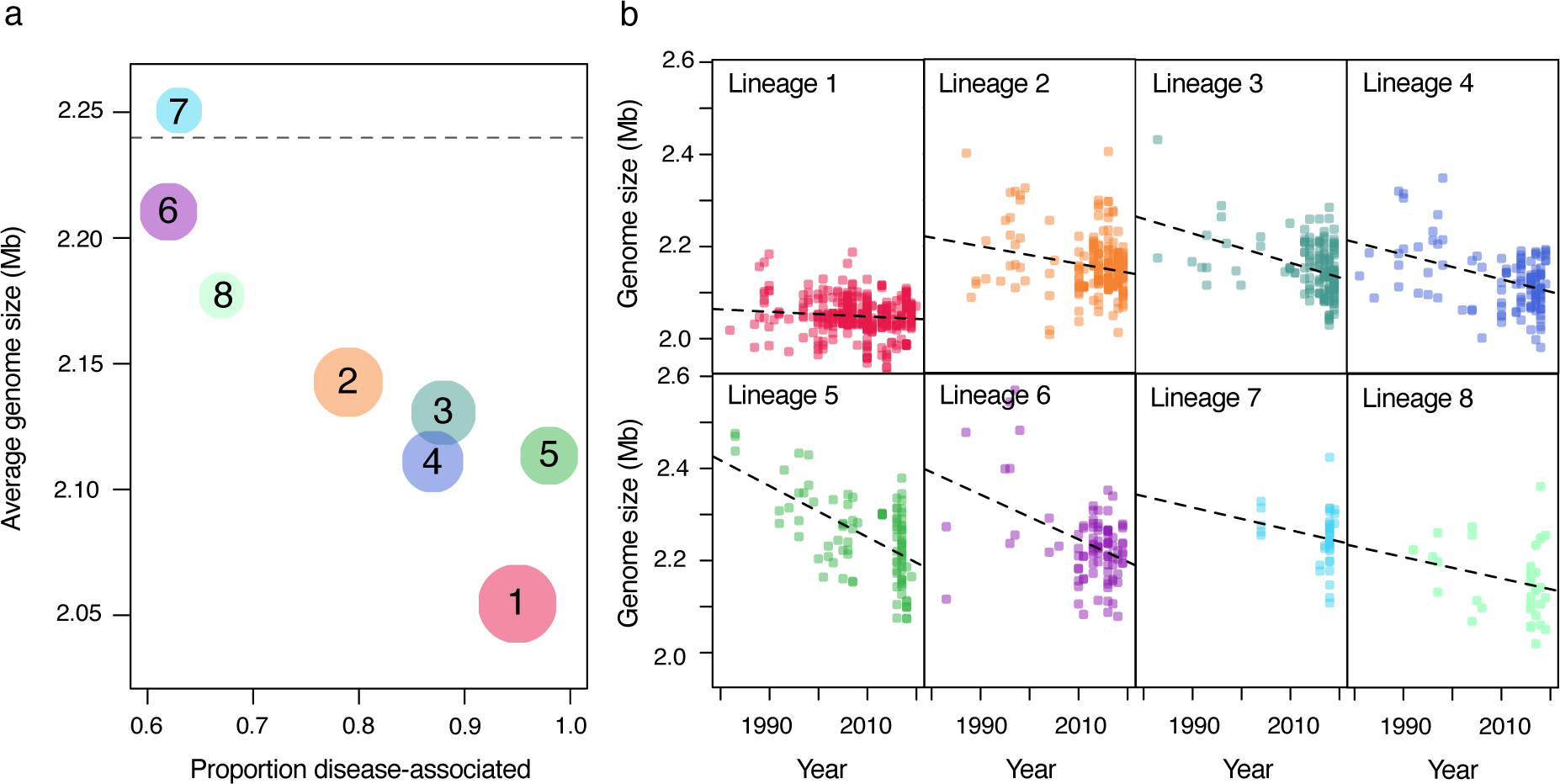
Gradual and ongoing genome reduction in pathogenic lineages. (a) Average genome sizes for recently sampled isolates (2018 to 2020) are shown against the proportion of disease-associated isolates for pathogenic lineages 1 to 8. Pathogenic lineages 9 and 10 are not shown as we have no or very few recent isolates from these lineages in our collection. The size of the points reflects the number of isolates in our collection on a log-scale. The dashed line represents the average genome size of isolates of *S. suis* from outside of the pathogenic clade, where disease isolates are present at a frequency of approximately 25%. (b) Genome sizes of individual isolates against sampling year for lineages 1 to 8. Dashed lines represent best fits for linear models of genome size against year of isolation; all have a negative slope, with a gradient ranging from a loss of 516 bases per year (lineage 1) to a loss of 5,550 bases per year (lineage 5).

Against this backdrop of genome reduction, we found evidence of adaptive gene acquisitions. In particular, we found evidence of multiple capsular switches in all of the six most common pathogenic lineages (Figure 1 and S24). Serotype 2 is often linked with invasive disease in pigs and zoonotic disease in humans, particularly when carried by ST1 (zoonotic lineage 1). We found that the serotype 2 capsule locus is present in the genomes of the majority of isolates from lineage 1 and also in isolates from 3/9 of the other pathogenic lineages (lineages 2, 4 and 9) and a few other lineages within the pathogenic clade (Table S1). While there is little diversity within the genes in the serotype 2 capsular locus either within or between lineages, there is greater diversity within zoonotic lineage 1 than within other lineages (Figure S25).

Combined with an older estimate of the most recent common ancestor of lineage 1 (1876, 95% CI: 1860-1891) compared to the clades that carry serotype 2 in lineages 2 (1935, 95% CI: 1923-1948) and 4 (1914, 95% CI: 1902-1926), this suggests that this serotype has been horizontally transferred from lineage 1 to these other pathogenic lineages. We also observe evidence of repeated transitions between serotypes 2 and 1/2 (whose capsular loci are nearly identical (34)). These transitions appear to be particularly common in lineage 2 (Figures S24 and S25).

Serotypes 1 and 14 are also common in zoonotic lineage 1. The genes encoding these capsules are largely shared with serotypes 2 and 1/2 (34), but divergence within the capsular genes is large enough to indicate independent origins of these two pairs of serotypes (Figure S24). Serotype 14 is also present in lineage 2, and in two other lineages (lineages 11 and 78). Diversity within the capsular genes of serotypes 1 and 14 is again consistent with them having been horizontally transmitted from zoonotic lineage 1 to other lineages, and repeated transitions between serotypes 1 and 14.

## Discussion

Studies of the impact of agricultural intensification on the risk of pathogen emergence generally focus on the most typical route: a pathogen jumping from one host species to another (4, 7, 35). In this study we instead described the emergence of an important zoonotic pathogen from within a largely commensal member of the respiratory microbiota of pigs. Our results suggest that while this form of pathogen emergence is gradual it can lead to a diverse pathogen whose impact on its host population is difficult to control, particularly alongside efforts to reduce the use of antibiotics in farming.

Over the last two hundred years global pig numbers have increased more than 10-fold (31). While the global rate of increase was highest in the second half of the 20^th^ century, in some regions, such as the United States, growth was faster in the 19^th^ century (Figure S15). Population growth has been accompanied by increased population density. For example, between 1921 and 2021 the number of pigs in Canada increased from 3.3 to 14.6 million, while the number of farms declined from 453 to just over 7 thousand (36). Modern farming practices also involve frequent movement of pigs between farms, with a trend towards production specialisation meaning that piglets often move to a different farm after weaning, and breeding for genetic improvement leading to the transport of breeding pigs around the world. These kinds of changes are predicted to promote pathogen emergence by facilitating the transmission of pathogens between hosts and therefore reducing the selective cost associated with increased host morbidity (8, 37).

Our analyses reveal high levels of genetic diversity in *S. suis* in pigs, that is mirrored in isolates sampled from a wild boar population in Spain. This diversity might reflect a long-standing association between *S. suis* and pigs (both domestic and wild). Our dating of the most recent common ancestors of the six most common pathogenic lineages in our collection indicates that they all emerged in the 19^th^ and 20^th^ centuries. The conclusion that these dates reflect an ecological shift towards pathogenicity in at least some of these lineages is supported by evidence that they coincided with the acquisition of a pathogenicity-associated genomic island (Island 3). It is further supported by patterns of genome reduction in each of the pathogenic lineages. In comparisons across bacterial species, it has been shown that bacterial pathogenicity is broadly associated with smaller genomes and fewer genes (22). While the drivers of this pattern are not well understood, and may be multiple, our observation of a gradual decline in the genome sizes of isolates from pathogenic lineages over our sampling period suggests a recent transition whose effect has yet to stabilise. Further evidence of an impact of intensive farming practices on pathogenic lineages of *S. suis* is found in our estimates of frequent long-distance movements, which are most likely to be a consequence of international trade in live animals.

While our results suggest that pathogenic lineages have emerged and spread globally over the last two centuries, they are also consistent with pathogenicity in *S. suis* predating this. Genetic diversity in all three of the pathogenicity-associated genomic islands we identified is consistent with their presence in *S. suis* long before the origins of individual pathogenic lineages. Genetic distances between the versions of Islands 1, 2 and 3 that are carried by lineages 1 and 2, are similar to distances between these lineages estimated from core genes. This may reflect the presence of these islands in a common ancestor of these two lineages. The prior existence of pathogenic lineages could therefore have aided the recent emergence of several new pathogenic lineages of *S. suis*.

Traits that promote virulence are thought to be selected for because they increase within-host growth or between-host transmission (38). The pathogenicity-associated genomic islands we identified have putative functions that may influence patterns of within-host growth. Islands 1 and 3 both have putative functions linked to metabolism, either the capacity to exploit particular sources of sugar within a host, or to regulate their metabolism. Metabolic capacity and growth rate have been linked to virulence in several bacterial species (39). The maintenance of both commensal and pathogenic lineages of *S. suis* could therefore be a consequence of a partitioning of the within-host niche. Pathogenic lineages may be better able to exploit particular regions of the tonsil than commensal lineages and vice versa, thereby reducing within-host competition. This could lead to segregation of these populations and reduced gene flow between them, which could in turn lead to the genome reduction in more pathogenic lineages due to fewer opportunities for gene acquisition from more diverse commensal lineages. This partitioning of the within-host niche may also be aided by Island 2. Pili are often involved in adhesion and evasion of cells, and therefore this island might aid in the colonisation of a particular region of the tonsil.

As we observe a high rate of spread of pathogenic lineages between countries, and previous studies have found that pathogenic strains of *S. suis* are capable of spreading rapidly between pigs within a farm (40), it is likely that pathogenic lineages have faster rates of between-host transmission than commensal lineages. As none of the pathogenicity-associated genes we identify have putative functions that are likely to be directly associated with between-host transmission, faster transmission rates may instead be driven by a trait with a diverse molecular basis, such as the capsular polysaccharide. In the human pathogen *Streptococcus pneumoniae* the capsular structure is known to both protect against immune recognition and immune clearance within a host (41), and promote between-host transmission through aiding survival outside a host (42), and increasing shedding (43).

The pathogenic lineages we have identified carry a mosaic of genes shared with other pathogenic and commensal lineages of *S. suis*. This is accompanied by diversity at key loci, such as the capsular locus, and in their association with systemic versus respiratory disease. Our results reveal that the emergence of novel pathogenic lineages therefore has not only led to more pathogenic lineages, but also to their diversification. This diversification of pathogenic lineages has led to difficulties in developing a cross-protective *S. suis* vaccine (44). We also find evidence of horizontal gene transfer between lineages leading to further diversification. In particular, we find evidence that the serotype 2 and serotype 1/14 capsules have been transferred from zoonotic lineage 1 (ST1) to other lineages. While we have been unable to explore this with our collection, which includes only a small sample of isolates from human infections, previous studies have suggested that strains of *S. suis* vary in their capacity to infect humans, and most human infections are caused by strains with a serotype 2 or 14 capsule (19). These acquisitions of the serotype 2 and 14 capsular loci may therefore have increased the zoonotic potential of these lineages.

Our results provide a new framework for understanding the genomic diversity in *S. suis* and its association with pathogenicity. This is likely to be of widespread use in *S. suis* research and in informing strategies for controlling the burden of this disease on pig farming and human health. As our collection spans only a small proportion of the countries that farm pigs globally, further sampling from a broader range countries and more extensive sampling within countries, particularly those with large and growing pig populations, is needed to investigate the existence of additional pathogenic lineages that are geographically restricted or have recently emerged. While our analyses suggest that the global movement of *S. suis* is driven by trade, they also suggest a possible role for wild boar. Further sampling of wild boar populations is needed to determine the role of wild boar in the transmission of pathogenic lineages between pig farms and into human populations. Further research is also required to experimentally characterise the functions of the pathogenicity-associated genomic islands we have identified, and establish their relationship with pathogenicity. Our results suggest that they represent evolutionary constraints on the emergence of pathogenic lineages: the emergence of new pathogenic lineages is contingent on the horizontal transfer of genes from an existing pathogenic lineage to another susceptible lineage. The conditions that allow for the spread of pathogenic lineages may therefore also promote the emergence of new pathogenic lineages and the diversification of existing pathogenic lineages through generating opportunities for the transfer of pathogenicity-associated genes. While this process has so far been gradual, we might predict that the pace will increase. Pig populations continue to grow, and the common pathogenic lineages we have identified have only recently spread to some parts of the world. This could lead to both an expanding niche for pathogenic lineages, and more opportunities for the emergence of new pathogenic lineages. Controlling the spread of pathogenic lineages of *S. suis* through pig populations should therefore be a priority to limit the potential impact of this pathogen on our future food security and public health.

## Methods

### Sampling of isolates and characterisation of disease-association

We generated a collection of genome assemblies of 3,070 isolates of *S. suis* and its close relatives including both newly sequenced isolates and previously published data (Table S1). The read data for newly sequenced isolates is publicly available from the SRA; the BioProject IDs are provided in Table S1. Our collection includes isolates from Australia, Canada, China, Denmark, Germany, Myanmar, the Netherlands, Spain, the UK, the USA and Vietnam. We also included 29 published reference genomes.

Isolates from Denmark, Germany, the Netherlands and Spain are largely from two new collections. The first aimed at collecting similar numbers of systemic disease, respiratory disease and carriage isolates from pigs from each country and were sampled from 2014 to 2018 (n=593). The second was a sample of historic isolates that aimed at capturing the breadth of disease-associated strains from each country from sample archives; they date from 1960 to 2007 (n=99). Isolates from the Netherlands also included a published collection of systemic disease isolates from humans and pigs that date from 1982 to 2008 (n=97) (45). Isolates from Spain also included newly sequenced isolates from a herd of wild boars sampled in 2015 as part of a published study (n=41) (46).

Isolates from the USA are from a newly sequenced collection of isolates from pigs from 2017 to 2020. Isolates were obtained from clinical cases submitted to the Iowa State University Veterinary Diagnostic Laboratory (ISU VDL) for routine diagnostics. Isolates from Myanmar are from another newly sequenced collection of isolates from pigs from farms and slaughterhouses in Yangon. They were sampled from pig farms that included small backyard farms, small-scale traditional farms, and modern industrial farms between 2016 and 2019. They were predominantly from throat swabs from pigs without *S. suis* disease, but were also sampled from farm and slaughterhouse drainage systems.

Isolates from the UK came from four different collections. The first sampled non-clinical and clinical isolates from pigs across England and Wales in 2009–2011 (described in (47)) (n=193). The second in 2013–2014 sampled non-clinical isolates from five farms (described in (48)) (n=117). The third in 2013–2014 targeted clinical isolates from pigs across England and Wales (described in (49)) (n=129). The fourth is a new collection of archived clinical isolates from pigs that date from 1987 to 2000 (n=49).

Isolates from Vietnam are from a collection that aimed at sampling closely related populations from humans and pigs (described in (47)). These included systemic disease isolates (n = 153) from human clinical cases of meningitis from provinces in southern and central Vietnam, and systemic disease (n = 6) or non-clinical isolates (n = 32) from pigs, collected between 2000 and 2010. These isolates were exclusively serotype 2 or 14. Isolates from Canada are from a previously published collection that aimed to target similar numbers of clinical and non-clinical isolates and dates from 1983 to 2016 (described in (50)).

Modern isolates from pigs from Denmark, Germany, the Netherlands, Canada, the UK and Vietnam were characterised as associated with systemic or respiratory disease, or non-clinical carriage based on clinical symptoms and isolation site. In pigs that showed clinical symptoms consistent with *S. suis* infections (e.g. meningitis, septicaemia and arthritis), the site of isolation was classified as ‘systemic’ if recovered from systemic sites (i.e. brain, liver, blood, joints). The site of recovery was classified as ‘respiratory’ if derived from lungs with gross lesions of pneumonia. *S. suis* isolates from the tonsils, nose or tracheo-bronchus of healthy pigs or dead pigs without any typical signs of *S. suis* infections were defined as ‘non-clinical’. Isolates that could not confidently be assigned to these categories (e.g. a tonsil isolate from a pig with systemic signs) were classified as unknown.

### Whole genome sequencing and assembly

Illumina whole genome sequencing was undertaken for all newly sequenced isolates. For newly sequenced isolates from Europe and the UK, DNA extraction, library preparation and sequencing was undertaken using a HiSeq 2500 instrument (Illumina, San Diego, CA, USA) by MicrobesNG (Birmingham, UK). For newly sequenced isolates from the USA, multiplex genome libraries were prepared using the Nextera XT DNA library preparation kit (Illumina, San Diego, CA, USA). The genomic library was quantified using a Qubit fluorometer dsDNA HS kit (Life Technologies Carlsbad, CA, USA) and normalized to the recommended amplification concentrations. The pooled libraries were sequenced on an Illumina Miseq sequencer using Miseq Reagent V3 for 600 cycles (Illumina, San Diego, CA, USA). Raw reads were demultiplexed automatically on the Miseq.

Raw sequence reads were pre-processed using Trimmomatic V0.36 to remove adaptors, trim poor quality ends, and delete short sequences (<36nt) (51). Raw and pre-processed reads were assessed for quality to ensure cleaning efficiency using FastQC (http://www.bioinformatics.babraham.ac.uk/projects/fastqc). We generated *de novo* assemblies with Spades v.3.12.0 (52). All assemblies were evaluated using QUAST v.5.0.1 (53) and we mapped reads back to *de novo* assemblies to investigate polymorphism (indicative of mixed cultures) using Bowtie2 v1.2.2 (54). Low quality assemblies were excluded from the collection.

We assembled high quality reference genomes for 19 isolates. For 12/19 isolates long-read sequencing library preparation was performed using Genomic-tips and a Genomic Blood and Cell Culture DNA Midi kit (Qiagen, Hilden, Germany). Sequencing was performed on the Sequel instrument from Pacific Biosciences using v2.1 chemistry and a multiplexed sample preparation. Reads were demultiplexed using Lima in the SMRT link software (https://github.com/PacificBiosciences/barcoding). Reads shorter than 2500 bases were removed using prinseq-lite.pl (https://sourceforge.net/projects/prinseq). Hybrid assemblies, using filtered PacBio and Illumina reads, and preliminary assemblies of long-read data assembled with Canu v1.91. (55) were generated with Unicycler v0.4.7 using the normal mode and default settings (56). Assembly graphs were visualised and, if necessary, manually corrected with Bandage (57). For the remaining 7/19 isolates, library preparation, short-read and long-read sequencing, and hybrid assembly with Unicycler were undertaken by MicrobesNG as part of their Enhanced Genome Service, which uses both Illumina and Oxford Nanopore Technologies. Four of these complete assemblies were published in a previous study (58).

### Serotyping and sequence-typing

Serotypes and sequence types were determined *in silico* using the Athey *et al.* (59) pipeline.

### Identification of homologous genes and analysis of pathogenicity-associated genomic islands

All genomes were annotated using Prokka v1.14.5 (60). Panaroo v1.2.2 (61) was used to identify orthologous genes (using recommended parameter settings) and create alignments of core genes. Pathogenicity-associated genomic islands were identified through comparison of frequencies of homologous genes identified by Panaroo and analysis of their relative genomic locations. Concatenated alignments of genes within each genomic island were generated and checked by eye. Regions too divergent to align were excluded from further analysis. Distance matrices were estimated and a neighbour-joining tree created using the *ape* package in R (62). Trees were visualised using iTOL and Grapetree (63, 64).

### Analysis of population structure, phylogeny and geographic spread

We used PopPunk to cluster our genomes (65). We first used the software to identify divergent genomes and then used the pipeline to cluster genomes in the central population into lineages. We used the standard pipeline followed by refinement using only core distances to determine lineages.

We generated reference-mapped assemblies of the six most common pathogenic lineages with Bowtie2 using reference genomes within the lineages. We identified recombination using Gubbins v2.3.1 (66) and masked all identified recombinant sites in our alignments. We tested for temporal signal in each lineage using a regression of root-to-tip distances against sampling year using the trees output from Gubbins and a published *R* script (67). Roots were chosen so as to minimise the residual mean squares of a linear regression. For lineage 1 we down sampled our data from 530 to 200 isolates that were randomly selected under the constraint of maintaining the temporal and geographic breadth of the collection. For each lineage we tested for temporal signal using 1,000 random permutations of dates over clades sampled from the same year to account for any confounding of temporal and genetic structure. This yielded significant evidence (*p* < 0.05) of temporal signal in all lineages except for in lineage 2. Nevertheless, as our estimate of evolutionary rate for lineage 2 was similar to lineages 1 and 3, we considered it likely that there was sufficient temporal signal to inform our estimates of dates.

We constructed dated phylogenies using BEAST v1.10 with a HKY+Γ model, a strict molecular clock, and an exponential population size coalescent model (68). We also constructed dated phylogenies using a relaxed molecular clock, a constant population size model, and a skyline population model, and found that our results were robust to different model choices. We undertook ancestral state reconstruction in BEAST to infer the geographic spread of these lineages by fitting an asymmetric discrete traits model to the posterior distributions of trees, with each state representing a country.

## Acknowledgements

This work was primarily funded by an EU Horizon 2020 grant “PIGSs” (727966) and a ZELS BBSRC award “Myanmar Pigs Partnership (MPP)” (BB/L018934/1). LAW, GGRM and EM was supported by a Sir Henry Dale Fellowship to LAW jointly funded by the Wellcome Trust and the Royal Society (109385/Z/15/Z). NH was supported by a Challenge grant from the Royal Society (CH16011) and an Isaac Newton Trust Research Grant (17.24(u)). GGRM was also supported by a Research Fellowship at Newnham College. SB is supported by the Medical Research Council (MR/V032836/1). PIC North America provided part of the funds for the sequencing of the isolates from the USA. MM and AJB were funded by Medical Research Council and Biotechnology and Biological Sciences Research Council studentships respectively, and MM was co-funded by the Raymond and Beverly Sackler Fund. We would like to acknowledge Susanna Williamson at the APHA for providing samples, Oscar Cabezón for sampling of the wild boar population in Spain, Mark O’Dea for access to sequence data from Australian isolates, the PIGSs and MPP consortiums for providing samples and helpful discussions, Julian Parkhill and John Welch for helpful discussions, and two anonymous reviewers for their valuable suggestions for improving the manuscript.

For the purpose of open access, we have applied a CC BY public copyright license to any author-accepted manuscript version arising from this submission.

