## Supplementary Figures and Table Legends for "The emergence and diversification of a zoonotic pathogen from within the microbiota of intensively farmed pigs"

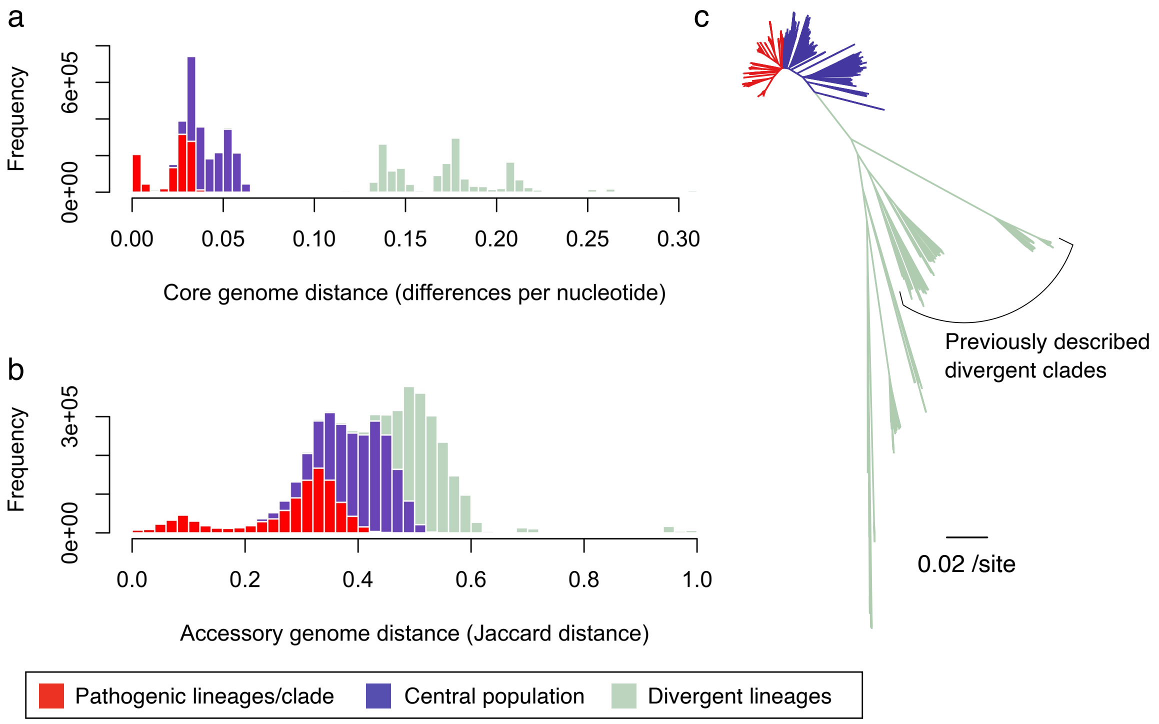


**Figure S1. Core and accessory genome distances across our collection of 3,070 isolates.** (a) Histogram of cophenetic pairwise distances estimated using the *ape* package in *R* based on a neighbour-joining tree constructed from a core gene alignment generated by *Panaroo*. (b) Histogram of Jaccard distances generated from a description of the presence/absence of homologous genes (excluding singletons) identified using *Panaroo*. (c) Phylogeny constructed from a concatenated core gene alignment constructed using the core gene alignment generated by *Panaroo* and a neighbour-joining algorithm with the *ape* package in *R*. In (a) and (b) comparisons between isolates in the 10 pathogenic lineages are shown in red, comparisons between isolates in the central *S. suis* population (but not between isolates in the 10 pathogenic lineages) are shown in deep blue, and comparisons between isolates from divergent lineages and between isolates from the central population and divergent isolates are shown in pale green. In (c) similar colours are used to represent lineages within the clade that contains the 10 pathogenic lineages (the pathogenic clade; red), lineages with the remainder of the central population (blue) and divergent lineages (pale green). The three divergent clades of *S. suis* that were described in a previous study (21) are indicated by a black bracket. It is possible that accessory genome distances are overestimated for more distantly related isolates due to the use of a strict cut-off for identifying homologous genes.

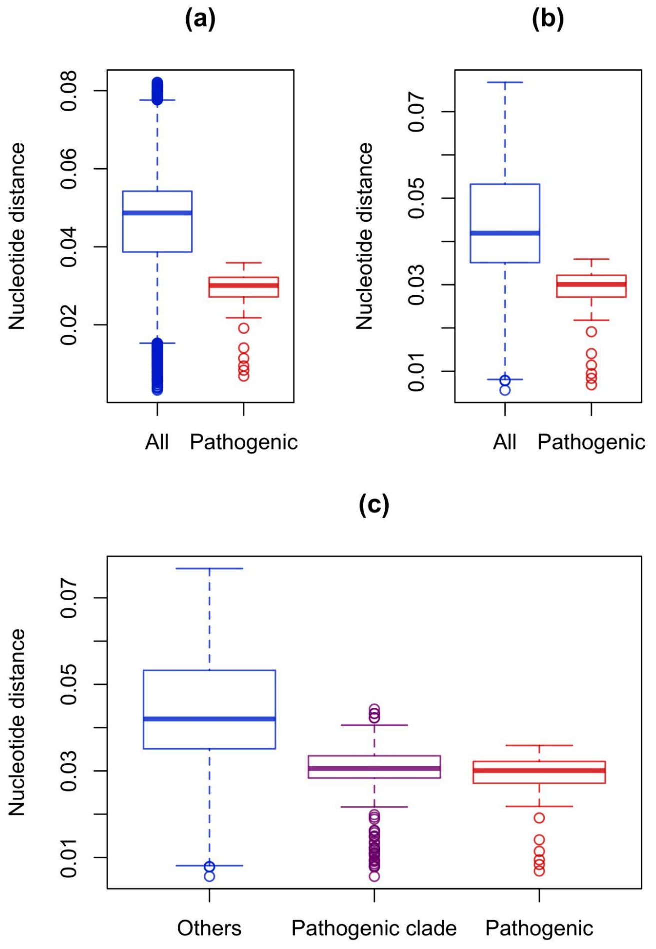


**Figure S2.** Pairwise core nucleotide distances between lineages within the central *S. suis* population. (a) Pairwise distances between all lineages we identify ("All", blue) and just between the 10 pathogenic lineages ("Pathogenic", red). (b) Pairwise distances between the 10 pathogenic lineages and other lineages in the central population ("All", blue) and just between the 10 pathogenic lineages ("Pathogen", red). (c) Pairwise distances between the 10 pathogenic lineages ("Pathogenic", red) compared to pairwise distances between the 10 pathogenic lineages and other 34 lineages in the clade in the core genome phylogeny that contains the 10 pathogenic lineages ("Pathogenic clade", purple), and all other lineages in the central *S. suis* population ("Others", blue).

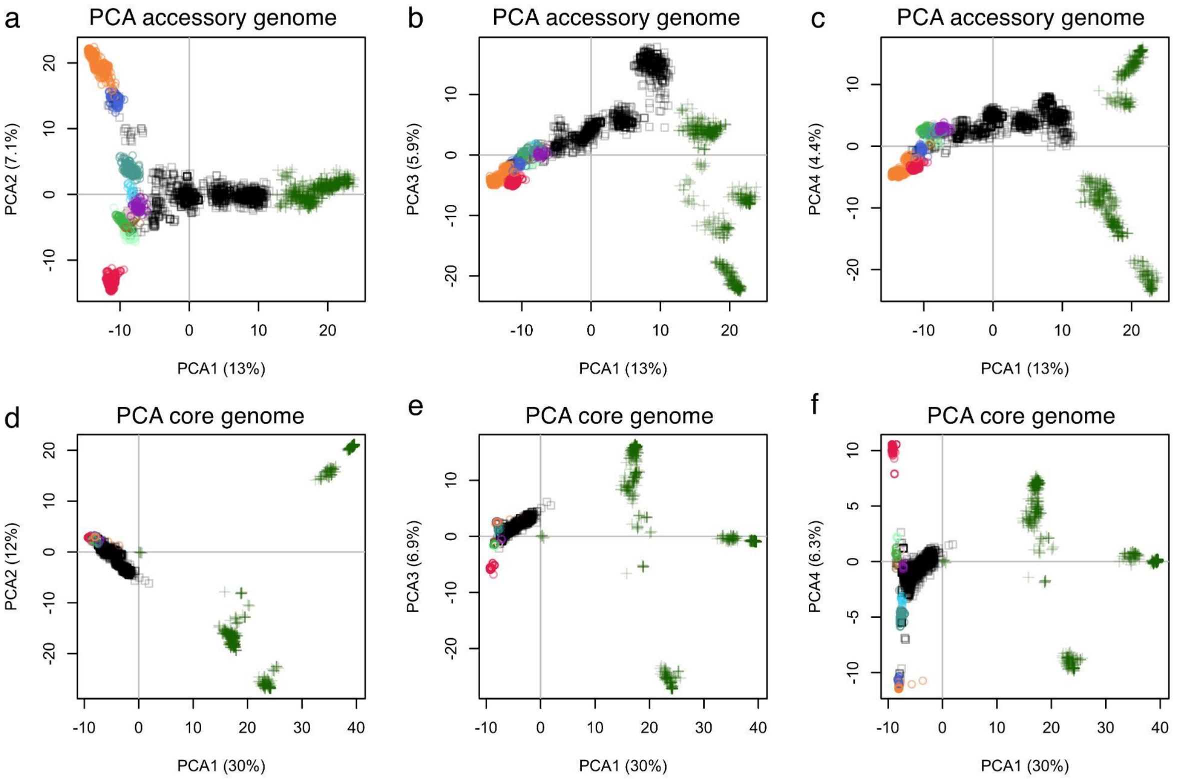


**Figure S3.** Principal component analysis (PCA) results for the accessory and core genomes. Isolates from the 10 pathogenic lineages are represented by circles coloured as in Figure 1, those from the central population outside of these lineages are represented by black squares, and those from divergent lineages green addition signs.

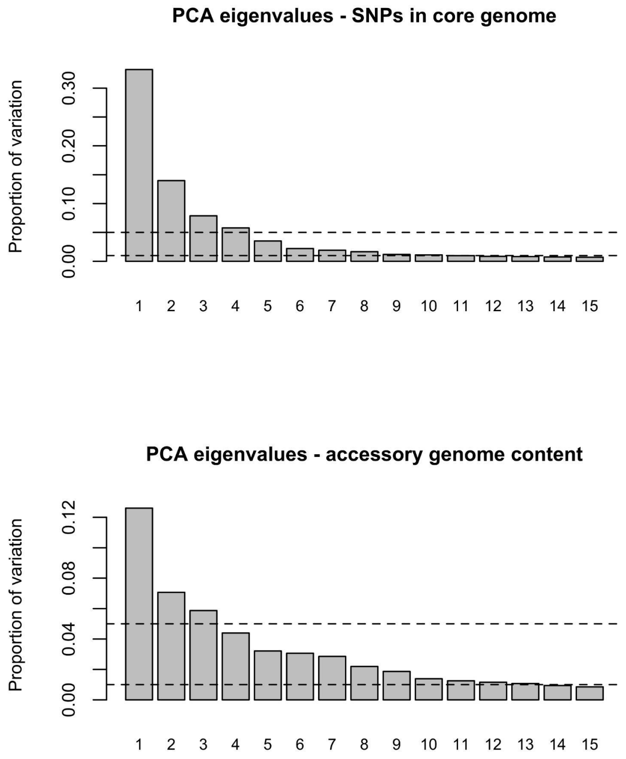


**Figure S4.** Principal component analysis eigenvalues for the accessory and core genomes.

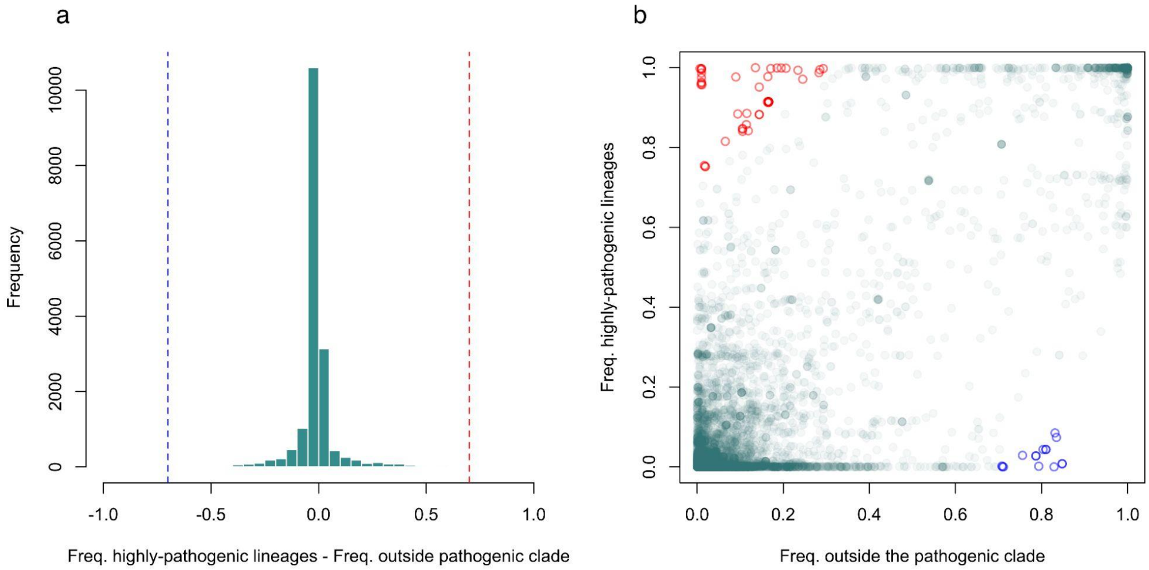


**Figure S5. Differences in gene frequency between isolates from the 10 pathogenic lineages and isolates from the central population outside of the pathogenic clade.** (a) Histogram of differences in gene frequency between the 1,523 isolates in the 10 pathogenic lineages and the 901 isolates from central population outside of the pathogenic clade. Cut-offs to identify genes that differ most in their frequency across these two groups are shown as red (>70% more common in pathogenic lineages) and blue (>70% more common outside of the pathogenic clade) dashed lines. (b) Frequencies of genes in set of isolates from the pathogenic lineages against their frequency in the set of isolates from the central population outside of the pathogenic clade. Genes that are >70% more frequent in isolates from pathogenic lineages are shown as red circles (top left) and those that are >70% more frequent in isolates from outside of the pathogenic clade are shown as blue circles (bottom right).

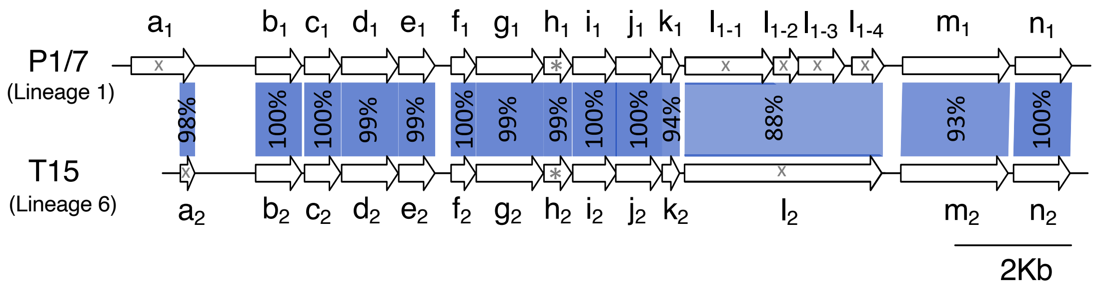


| Gene IDs | | Reference Genomes | | Frequency in pathogenic lineages | | | | | | | | | |
|---|---|---|---|---|---|---|---|---|---|---|---|---|---|
| | | P1/7 ID | T15 ID | 1 | 2 | 3 | 4 | 5 | 6 | 7 | 8 | 9 | 10 |
| $a_1$ | $a_2$ | SSU_RS05405 | T15_RS03830 | 1.00 | 1.00 | 0.30 | 1.00 | 1.00 | 1.00 | 1.00 | 1.00 | 1.00 | 1.00 |
| $b_1$ | $b_2$ | SSU_RS05400 | T15_RS03835 | 1.00 | 1.00 | 0.30 | 1.00 | 1.00 | 1.00 | 1.00 | 1.00 | 1.00 | 1.00 |
| $c_1$ | $c_2$ | SSU_RS05395 | T15_RS03840 | 1.00 | 1.00 | 0.30 | 1.00 | 1.00 | 1.00 | 1.00 | 1.00 | 1.00 | 1.00 |
| $d_1$ | $d_2$ | SSU_RS05390 | T15_RS03845 | 1.00 | 1.00 | 0.30 | 1.00 | 1.00 | 1.00 | 1.00 | 1.00 | 1.00 | 1.00 |
| $e_1$ | $e_2$ | SSU_RS05385 | T15_RS03850 | 1.00 | 1.00 | 0.30 | 1.00 | 1.00 | 1.00 | 1.00 | 1.00 | 1.00 | 1.00 |
| $f_1$ | $f_2$ | SSU_RS05380 | T15_RS03855 | 1.00 | 1.00 | 0.30 | 1.00 | 1.00 | 1.00 | 1.00 | 1.00 | 1.00 | 1.00 |
| $g_1$ | $g_2$ | SSU_RS05375 | T15_RS03860 | 1.00 | 1.00 | 0.30 | 1.00 | 1.00 | 1.00 | 1.00 | 1.00 | 1.00 | 1.00 |
| $h_1$ | $h_2$ | SSU_RS05370 | T15_RS03865 | 1.00 | 1.00 | 0.30 | 1.00 | 1.00 | 1.00 | 1.00 | 1.00 | 1.00 | 1.00 |
| $i_1$ | $i_2$ | SSU_RS05365 | T15_RS03870 | 1.00 | 1.00 | 0.30 | 1.00 | 1.00 | 1.00 | 1.00 | 1.00 | 1.00 | 1.00 |
| $j_1$ | $j_2$ | SSU_RS05360 | T15_RS03875 | 1.00 | 1.00 | 0.30 | 1.00 | 1.00 | 1.00 | 1.00 | 1.00 | 1.00 | 1.00 |
| $k_1$ | $k_2$ | SSU_RS05355 | T15_RS03880 | 1.00 | 1.00 | 0.30 | 1.00 | 1.00 | 1.00 | 1.00 | 1.00 | 1.00 | 1.00 |
| $l_{1-1}$; $l_{1-2}$; $l_{1-3}$; $l_{1-4}$ | $l_2$ | SSU_RS05335; SSU_RS05340; SSU_RS05345; SSU_RS05350 | T15_RS03885 | 1.00 | 0.99 | 0.30 | 1.00 | 1.00 | 1.00 | 1.00 | 1.00 | 1.00 | 1.00 |
| $m_1$ | $m_2$ | SSU_RS05330 | T15_RS03890 | 1.00 | 0.98 | 0.30 | 1.00 | 1.00 | 1.00 | 1.00 | 1.00 | 1.00 | 1.00 |
| $n_1$ | $n_2$ | SSU_RS05325 | T15_RS03895 | 1.00 | 1.00 | 0.30 | 1.00 | 1.00 | 1.00 | 1.00 | 1.00 | 1.00 | 1.00 |

| Gene IDs | | Annotation |
|---|---|---|
| $a_1$ | $a_2$ | Transposase |
| $b_1$ | $b_2$ | NAD(P)-dependent dehydrogenase, short-chain alcohol dehydrogenase family (lipid transport and metabolism, secondary metabolites biosynthesis, transport and catabolism) |
| $c_1$ | $c_2$ | Ribose 5-phosphate isomerase RpiB (carbohydrate transport and metabolism) |
| $d_1$ | $d_2$ | Sugar or nucleoside kinase, ribokinase family (carbohydrate transport and metabolism) |
| $e_1$ | $e_2$ | 2-keto-3-deoxy-6-phosphogluconate aldolase (carbohydrate transport and metabolism) |
| $f_1$ | $f_2$ | Phosphotransferase system, mannose/fructose/N-acetylgalactosamine-specific component IIA (carbohydrate transport and metabolism) |
| $g_1$ | $g_2$ | Unknown function |
| $h_1$ | $h_2$ | Phosphotransferase system, mannose/fructose/N-acetylgalactosamine-specific component IIB (carbohydrate transport and metabolism) |
| $i_1$ | $i_2$ | Phosphotransferase system, mannose/fructose/N-acetylgalactosamine-specific component IIC (carbohydrate transport and metabolism) |
| $j_1$ | $j_2$ | Phosphotransferase system, mannose/fructose/N-acetylgalactosamine-specific component IID (carbohydrate transport and metabolism) |
| $k_1$ | $k_2$ | Preprotein translocase subunit YajC (intracellular trafficking, secretion, and vesicular transport) |
| $l_{1-1}$; $l_{1-2}$; $l_{1-3}$; $l_{1-4}$ | $l_2$ | Unknown function |
| $m_1$ | $m_2$ | Heparinase |
| $n_1$ | $n_2$ | DNA-binding transcriptional regulator, LacI/PurR family (transcription) |

**Figure S6. Genes in Island 1.** The image shows examples of Island 1 from two reference genomes in lineages 1 and 6. Blue shading indicates homology; deeper colours represent greater protein sequence identity (percentage identity shown). * indicates the gene used to describe the presence/absence of the island (shown in Figure 2). x indicates genes excluded from our phylogenetic reconstructions (shaded grey in the two tables). The two tables provide gene IDs in the two reference strains, the proportion of isolates from each of the pathogenic lineages that these genes are found in, and an annotation based on an NCBI protein-level BLAST.

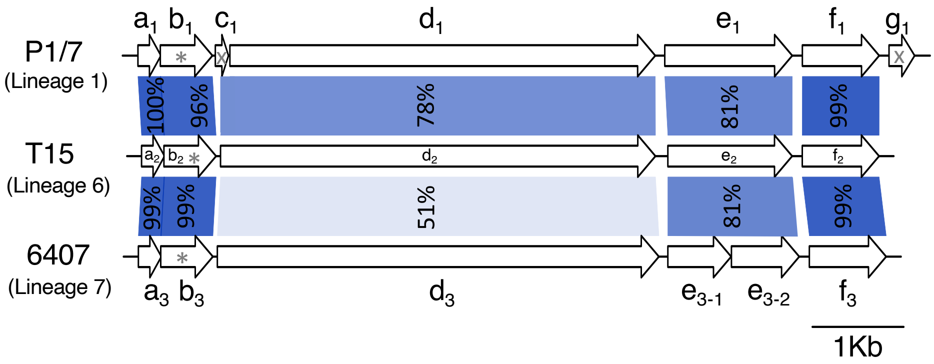


| Gene IDs | | | Reference Genomes | | | Frequency in pathogenic lineages | | | | | | | | | |
|---|---|---|---|---|---|---|---|---|---|---|---|---|---|---|---|
| | | | P1/7 ID | T15 ID | 6407 ID | 1 | 2 | 3 | 4 | 5 | 6 | 7 | 8 | 9 | 10 |
| $a_1$ | $a_2$ | $a_3$ | SSU_RS 02325 | T15_RS 02515 | ID09_RS 02305 | 1.00 | 0.82 | 0.97 | 1.00 | 0.01 | 1.00 | 1.00 | 0.03 | 0.00 | 1.00 |
| $b_1$ | $b_2$ | $b_3$ | SSU_RS 02330 | T15_RS 02520 | ID09_RS 02310 | 1.00 | 0.82 | 0.97 | 1.00 | 0.01 | 1.00 | 1.00 | 0.00 | 0.00 | 1.00 |
| $c_1$ | $d_2$ | $d_3$ | SSU_RS 02335 | T15_RS 02525 | ID09_RS 02315 | 1.00 | 0.82 | 0.97 | 1.00 | 0.01 | 1.00 | 1.00 | 0.00 | 0.00 | 1.00 |
| $d_1$ | | | SSU_RS 02340 | | | | | | | | | | | | |
| $e_1$ | $e_2$ | $e_{3\text{-}1}$; $e_{3\text{-}2}$ | SSU_RS 02345 | T15_RS 02530 | ID09_RS 02320 | 0.99 | 0.82 | 0.97 | 0.33 | 0.00 | 1.00 | 1.00 | 0.00 | 0.00 | 1.00 |
| $f_1$ | $f_2$ | $f_3$ | SSU_RS 02350 | T15_RS 02535 | ID09_RS 02325 | 0.99 | 0.82 | 0.95 | 0.99 | 0.00 | 0.98 | 1.00 | 0.00 | 0.00 | 1.00 |
| $g_1$ | - | - | SSU_RS 09995 | | | 1.00 | 0.78 | 0.01 | 0.77 | 0.01 | 0.20 | 0.05 | 0.03 | 0.00 | 0.00 |

| Gene IDs | | | Annotation |
|---|---|---|---|
| $a_1$ | $a_2$ | $a_3$ | Unknown function |
| $b_1$ | $b_2$ | $b_3$ | Signal peptidase I (intracellular trafficking, secretion, and vesicular transport) |
| $c_1$ | $d_2$ | $d_3$ | Minor pilin subunit |
| $d_1$ | | | |
| $e_1$ | $e_2$ | $e_{3\text{-}1}$; $e_{3\text{-}2}$ | Major pilin subunit |
| $f_1$ | $f_2$ | $f_3$ | Sortase (surface protein transpeptidase) (cell wall/membrane/envelope biogenesis) |
| $g_1$ | - | - | Transposase |

**Figure S6. Genes in Island 2.** The image shows examples of Island 2 from three reference genomes in lineages 1, 6 and 8. Blue shading indicates homology; deeper colours represent greater protein sequence identity (percentage identity shown). * indicates the gene used to describe the presence/absence of the island (shown in Figure 2). x indicates genes excluded from our phylogenetic reconstructions (shaded grey in the two tables). The two tables provide gene IDs in the two reference strains, the proportion of isolates from each of the pathogenic lineages that these genes are found in, and an annotation based on an NCBI protein-level BLAST.

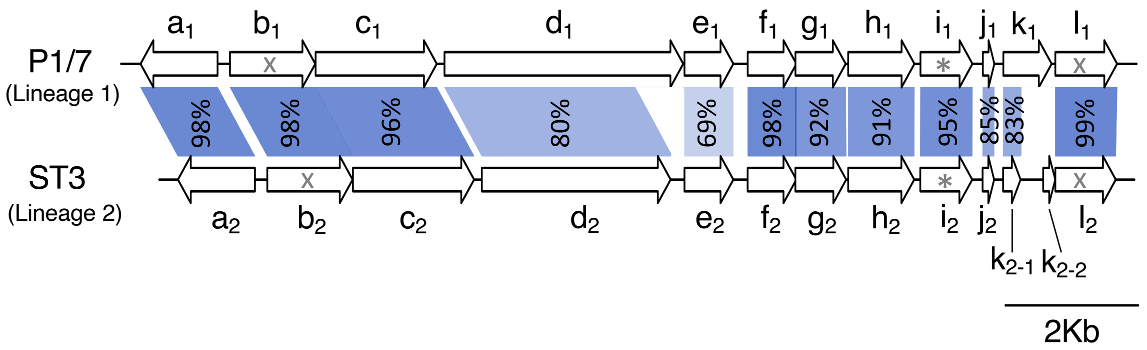


| Gene IDs | | Reference Genomes | | Frequency in pathogenic lineages | | | | | | | | | |
|---|---|---|---|---|---|---|---|---|---|---|---|---|---|
| | | **P1/7 ID** | **ST3 ID** | **1** | **2** | **3** | **4** | **5** | **6** | **7** | **8** | **9** | **10** |
| $a_1$ | $a_2$ | SSU_RS01130 | SSUST3_RS01160 | 0.98 | 0.94 | 0.88 | 0.96 | 1.00 | 0.95 | 0.97 | 1.00 | 0.96 | 1.00 |
| $b_1$ | $b_2$ | SSU_RS01135 | SSUST3_RS01165 | 1.00 | 1.00 | 0.99 | 0.99 | 1.00 | 0.99 | 0.97 | 1.00 | 0.96 | 1.00 |
| $c_1$ | $c_2$ | SSU_RS01140 | SSUST3_RS01170 | 1.00 | 1.00 | 0.99 | 0.99 | 1.00 | 0.99 | 0.95 | 1.00 | 0.96 | 1.00 |
| $d_1$ | $d_2$ | SSU_RS01145 | SSUST3_RS01175 | 0.99 | 0.99 | 0.97 | 1.00 | 1.00 | 0.98 | 1.00 | 0.97 | 0.96 | 1.00 |
| $e_1$ | $e_2$ | SSU_RS01150 | SSUST3_RS01180 | 0.97 | 0.98 | 0.95 | 0.98 | 0.96 | 0.96 | 1.00 | 0.76 | 0.96 | 1.00 |
| $f_1$ | $f_2$ | SSU_RS01155 | SSUST3_RS01185 | 0.99 | 0.86 | 0.99 | 1.00 | 0.92 | 0.98 | 1.00 | 0.79 | 0.96 | 1.00 |
| $g_1$ | $g_2$ | SSU_RS01160 | SSUST3_RS01190 | 1.00 | 0.86 | 0.99 | 1.00 | 0.96 | 0.97 | 1.00 | 0.79 | 0.96 | 1.00 |
| $h_1$ | $h_2$ | SSU_RS01165 | SSUST3_RS01195 | 1.00 | 0.97 | 0.93 | 0.99 | 0.98 | 0.98 | 1.00 | 0.76 | 0.96 | 1.00 |
| $i_1$ | $i_2$ | SSU_RS01170 | SSUST3_RS01200 | 1.00 | 1.00 | 0.99 | 1.00 | 1.00 | 0.99 | 1.00 | 1.00 | 0.96 | 1.00 |
| $j_1$ | $j_2$ | SSU_RS01175 | SSUST3_RS01205 | 1.00 | 1.00 | 0.99 | 1.00 | 1.00 | 0.98 | 1.00 | 1.00 | 0.96 | 1.00 |
| $k_1$ | $k_{2-1}$ | SSU_RS01180 | SSUST3_RS01210 | 1.00 | 1.00 | 0.99 | 1.00 | 1.00 | 0.99 | 1.00 | 1.00 | 0.96 | 1.00 |
| $l_1$ | $l_2$ | SSU_RS01185 | SSUST3_RS01215 | 0.98 | 0.98 | 0.96 | 0.97 | 1.00 | 0.94 | 1.00 | 1.00 | 0.96 | 1.00 |

| Gene IDs | | **Annotation** (based on NCBI BLAST) |
|---|---|---|
| $a_1$ | $a_2$ | ROK-family repressor protein (transcription, carbohydrate transport and metabolism) |
| $b_1$ | $b_2$ | Phosphotransferase system cellobiose-specific component IIC (carbohydrate transport and metabolism) |
| $c_1$ | $c_2$ | Unknown function |
| $d_1$ | $d_2$ | Unknown function |
| $e_1$ | $e_2$ | Unknown function |
| $f_1$ | $f_2$ | ABC-type nitrate/sulfonate/bicarbonate transport system, permease component (inorganic ion transport and metabolism) |
| $g_1$ | $g_2$ | ABC-type nitrate/sulfonate/bicarbonate transport system, ATPase component (inorganic ion transport and metabolism) |
| $h_1$ | $h_2$ | ABC-type nitrate/sulfonate/bicarbonate transport system, periplasmic component (inorganic ion transport and metabolism) |
| $i_1$ | $i_2$ | 6-phosphogluconolactonase/Glucosamine-6-phosphate isomerase/deaminase (carbohydrate transport and metabolism) |
| $j_1$ | $j_2$ | Cation transport ATPase (inorganic ion transport and metabolism) |
| $k_1$ | $k_{2-1}$ | Cation transport ATPase (inorganic ion transport and metabolism) |
| $l_1$ | $l_2$ | Cation transport ATPase (inorganic ion transport and metabolism) |

**Figure S8. Genes in Island 3.** The image shows examples of Island 3 from two reference genomes in lineages 1 and 2. Blue shading indicates homology; deeper colours represent greater protein sequence identity (percentage identity shown). * indicates the gene used to describe the presence/absence of the island (shown in Figure 2). x indicates genes excluded from our phylogenetic reconstructions (shaded grey in the two tables). The two tables provide gene IDs in the two reference strains, the proportion of isolates from each of the pathogenic lineages that these genes are found in, and an annotation based on an NCBI protein-level BLAST.

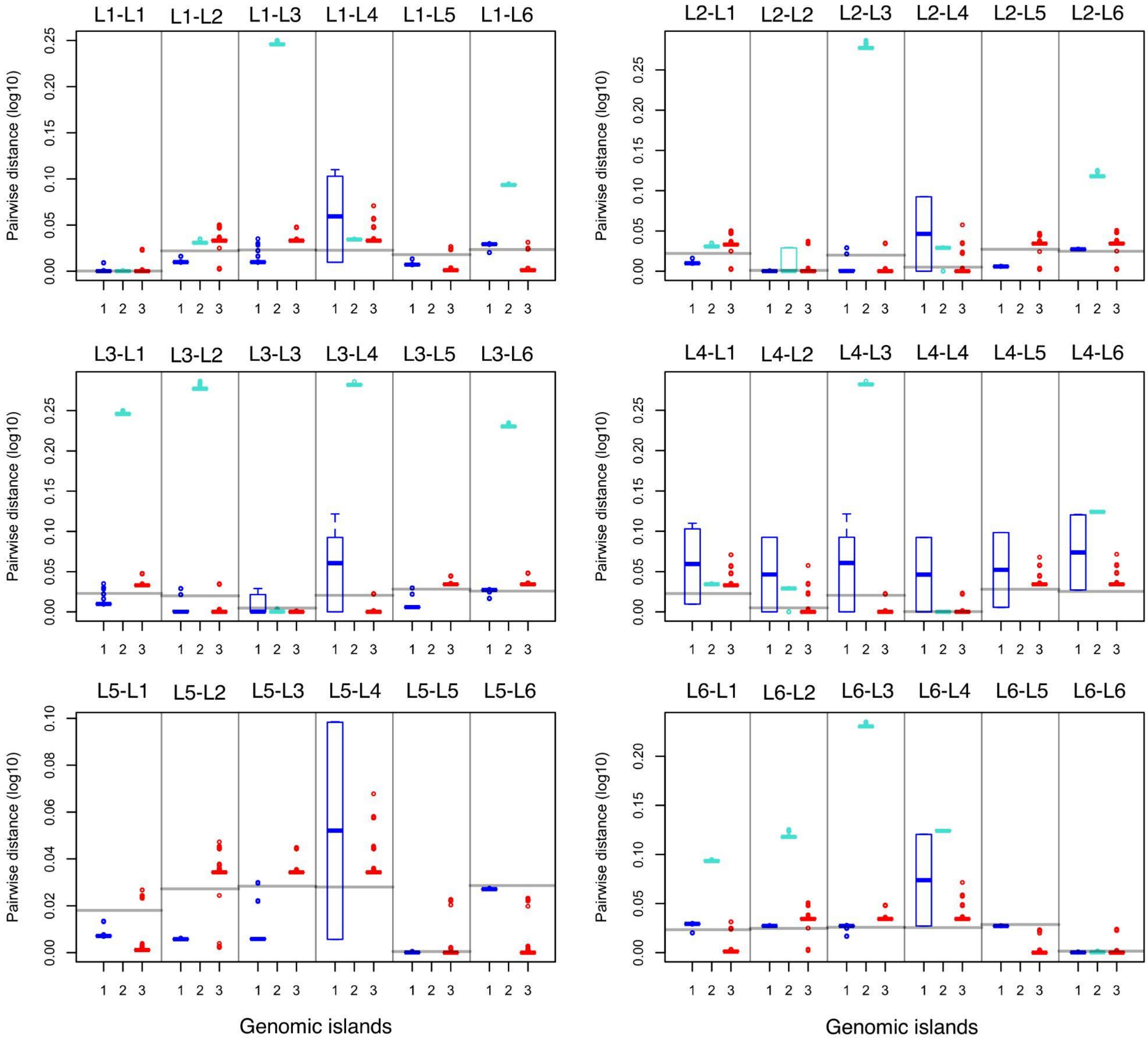


**Figure S9. Pairwise distances between isolates from the six most common pathogenic lineages for the 3 pathogenicity-associated genomic islands.** Boxplots are shown for each of the 3 genomic islands (labelled 1-3) with Island 1 in blue, Island 2 in turquoise and Island 3 in red. Grey horizontal lines represent the median values for core genes. Pairwise distances are shown within pathogenic lineages (e.g. L1-L1) and between pathogenic lineages (e.g. L1-L2). Pairwise distances are based on regions of islands that are shared and alignable across all isolates.

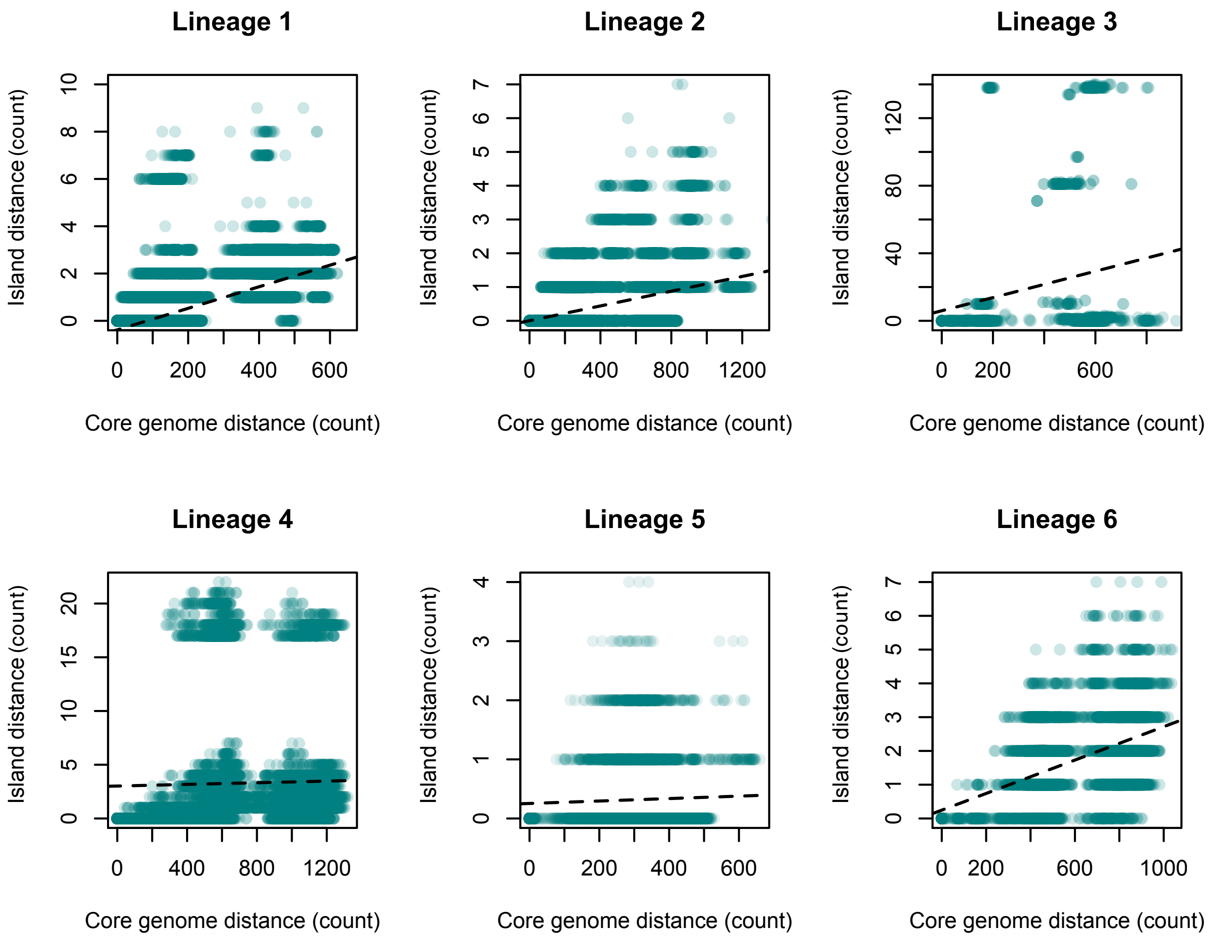


**Figure S10.** Comparisons of pairwise distances between isolates based on Island 1 and a recombination-stripped reference mapped alignment of the entire genome. Island distances are shown in counts of differences estimated directly from a nucleotide alignment while core genome distances are shown in counts of differences based on cophenetic distances from the recombination-stripped phylogeny that is produced by Gubbins. Dashed lines represent lines of best fit from a general linear regression. One isolate (M101513_S1) was excluded from our plot of Lineage 1 as it carries an unusually divergent version of the island (although it falls within the wider diversity present in other pathogenic lineages).

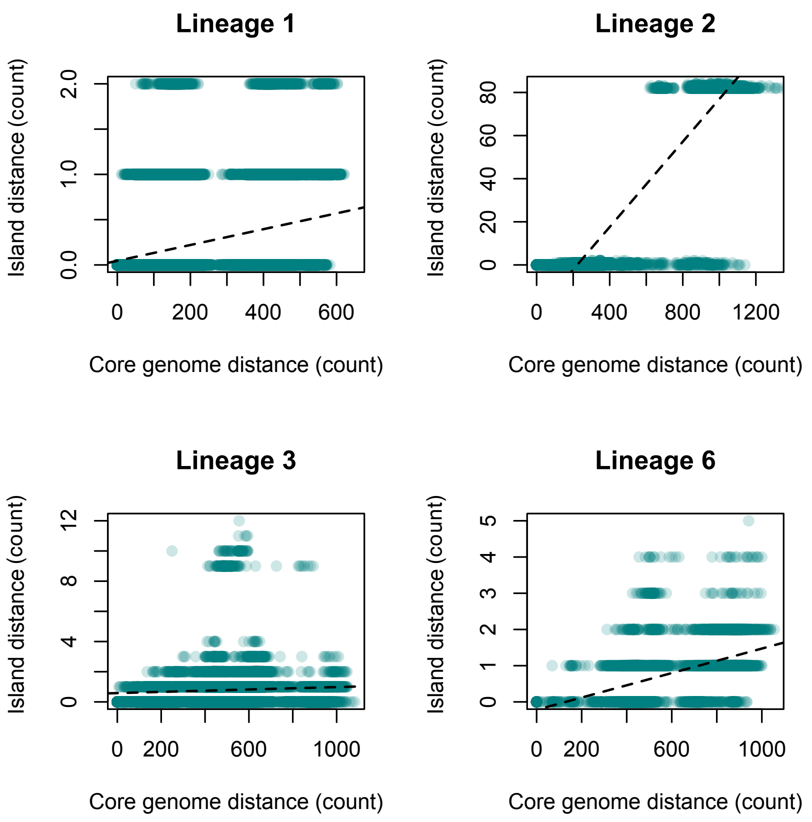


**Figure S11**. Comparisons of pairwise distances between isolates based on Island 2 and a recombination-stripped reference mapped alignment of the entire genome. Island distances are shown in counts of differences estimated directly from a nucleotide alignment while core genome distances are shown in counts of differences based on cophenetic distances from the recombination-stripped phylogeny that is produced by Gubbins. Dashed lines represent lines of best fit from a general linear regression.

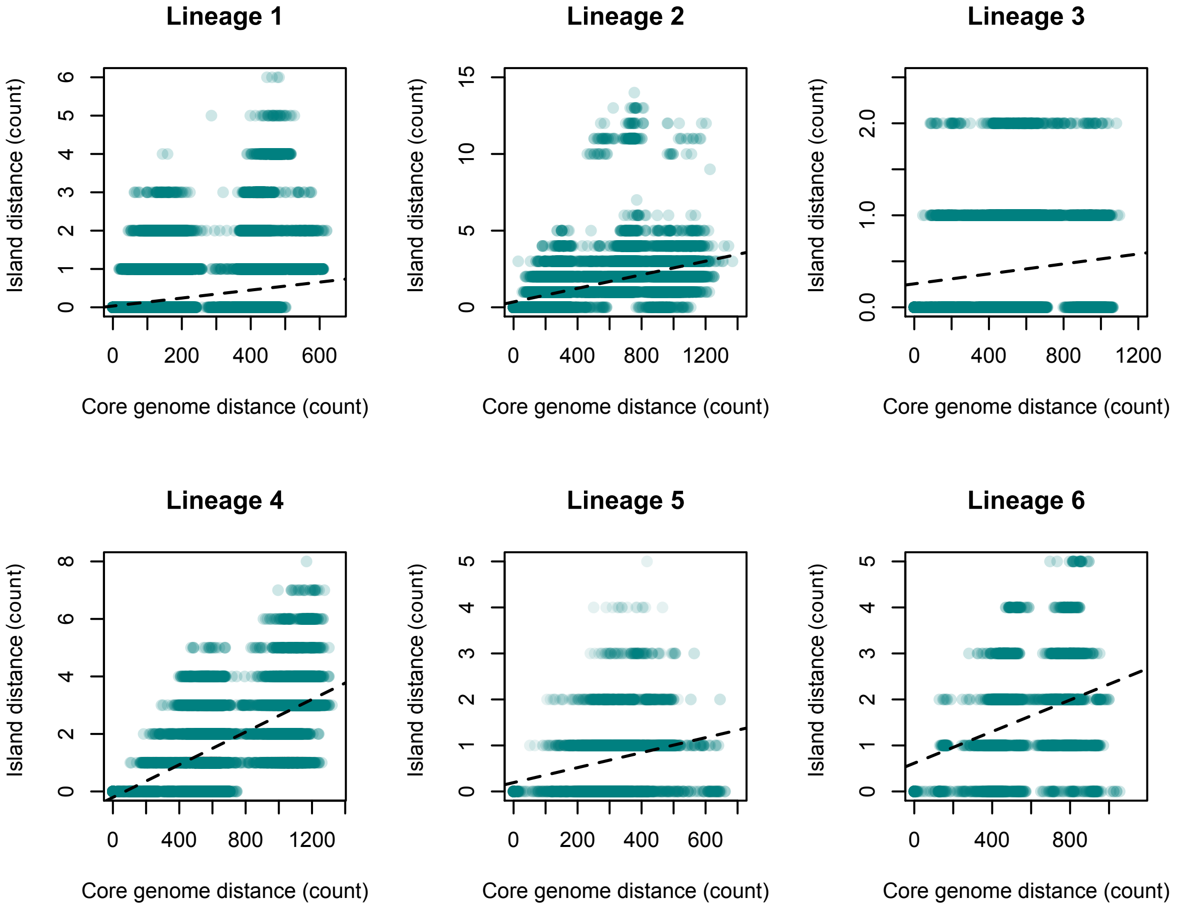


**Figure S12**. Comparisons of pairwise distances between isolates based on Island 3 and a recombination-stripped reference mapped alignment of the entire genome. Island distances are shown in counts of differences estimated directly from a nucleotide alignment while core genome distances are shown in counts of differences based on cophenetic distances from the recombination-stripped phylogeny that is produced by Gubbins. Dashed lines represent lines of best fit from a general linear regression.

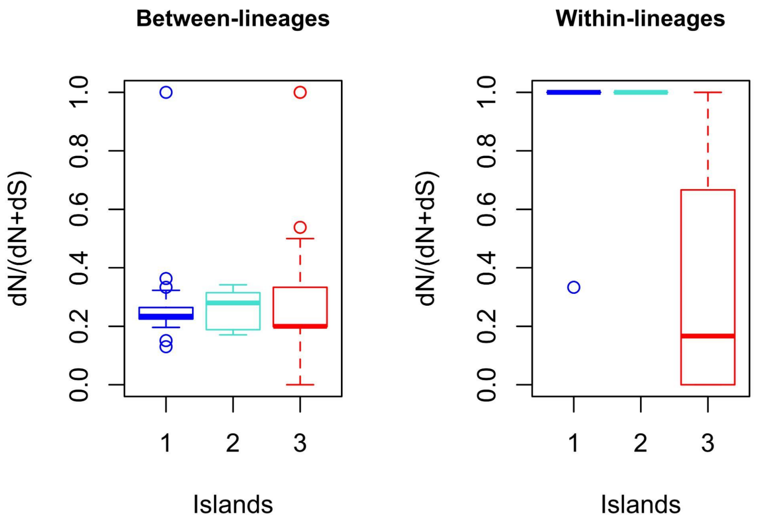


**Figure S13**. Relative pairwise distances at 1st and 2nd codon positions relative to 3rd codon positions for the three pathogenicity-associated islands (1, 2 and 3). Comparisons are shown for pairwise comparisons between-lineages and within-lineages.

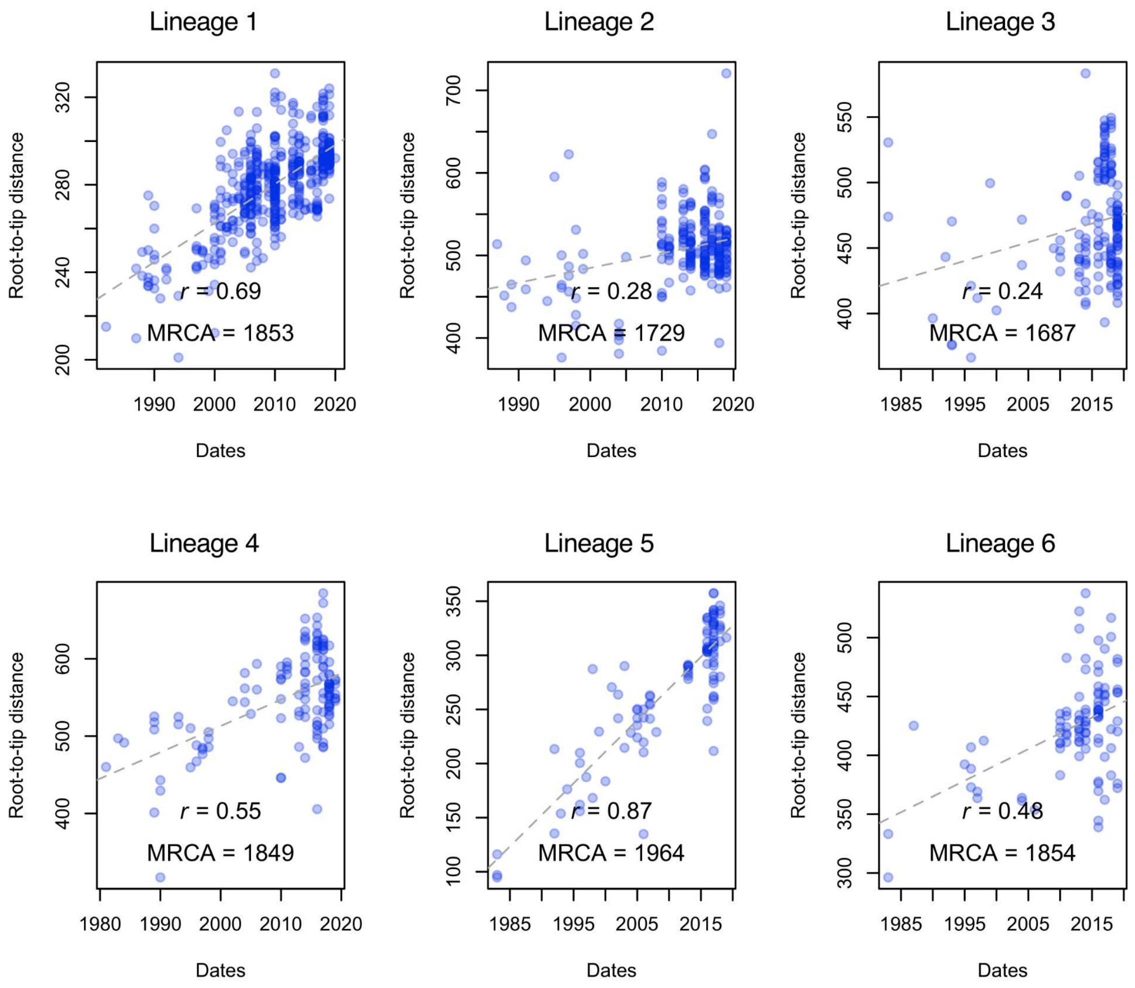


**Figure S14. Regressions of root-to-tip distances against sampling year for each of the six most common pathogenic lineages.** The phylogenies used for this analysis were the trees output from *Gubbins* following the stripping of recombinant sites from reference-mapped assemblies of each lineage. Roots were chosen so as to minimise the residual-mean-squares of a linear regression of root-to-tip distance against sampling date using a published *R* script [51] The dashed lines represent lines of best fit. The Pearson *r* and the value of the intercept (most recent common ancestor; MRCA) are reported for each lineage. 1000 random permutations were performed using clustering of clades sampled from the same year to account for any confounding of temporal and genetic structure for each lineage. All lineages except for lineage 2 showed significant evidence of temporal signal.

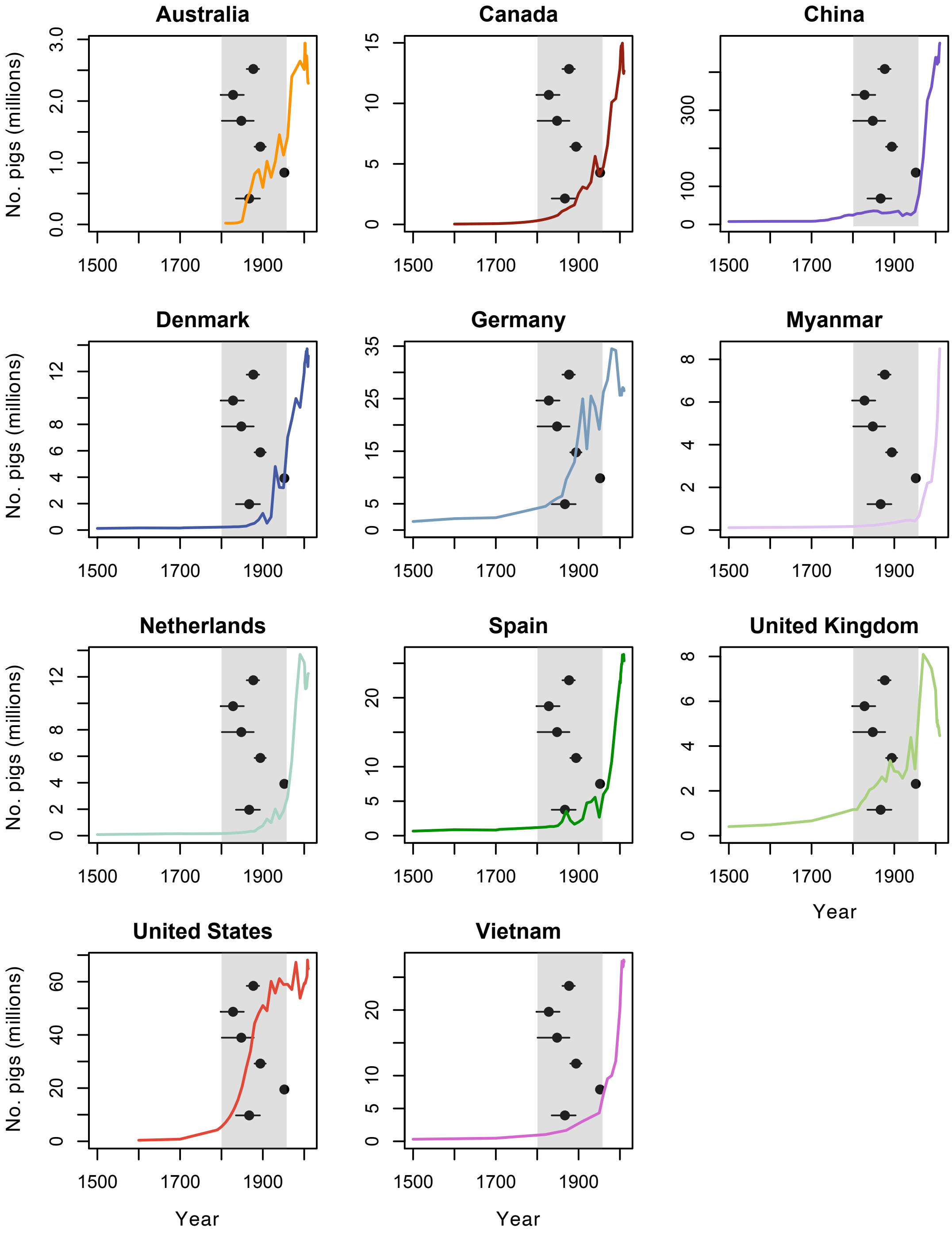


**Figure S15. Estimates of the dates of the most recent common ancestors of the six most common pathogenic lineages and country-specific estimates of historic pig numbers for each of the countries represented in our collection.** Grey shading represents the range of estimates for the origins of the six most common pathogenic lineages (including 95% confidence intervals).

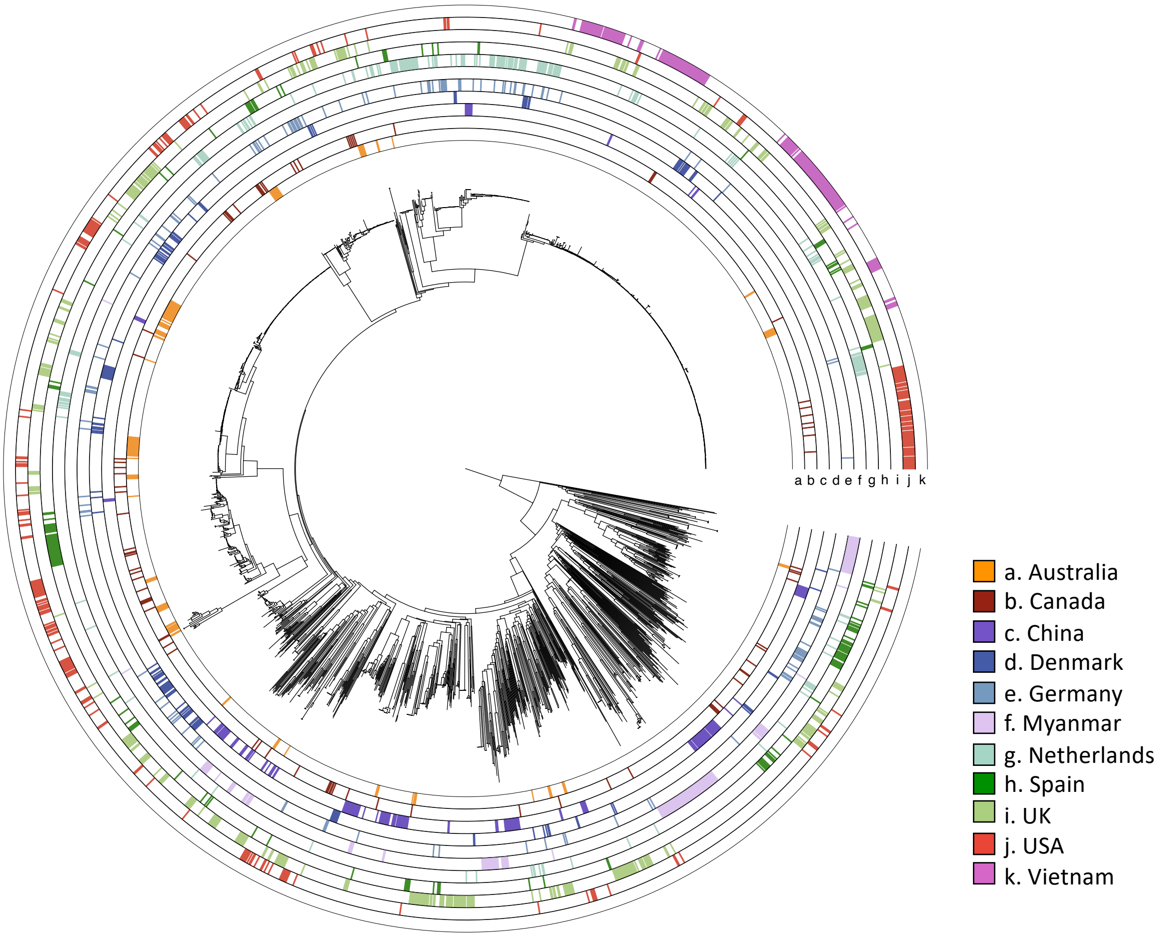


**Figure S16. Countries of origin mapped onto a core genome phylogeny of the central population of *S. suis*.**

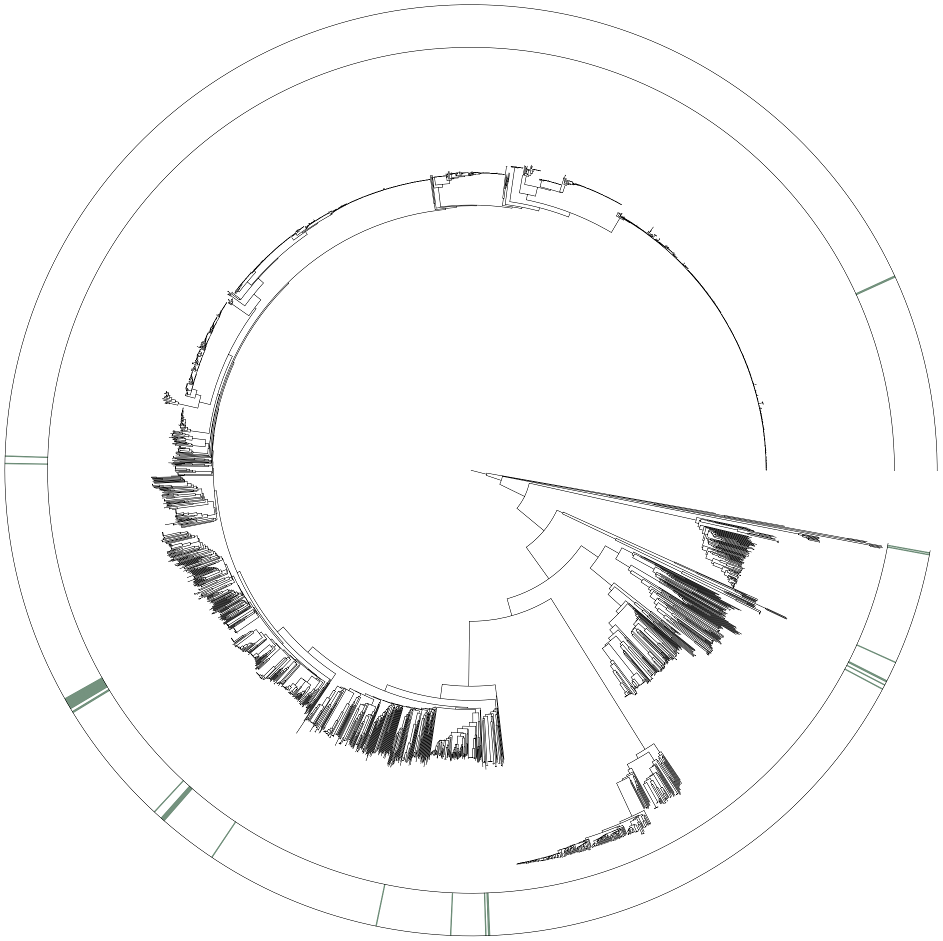

**Figure S17. Isolates from a wild boar population in Spain mapped onto a core genome phylogeny of the entire diversity present in our collection of *S. suis*.**

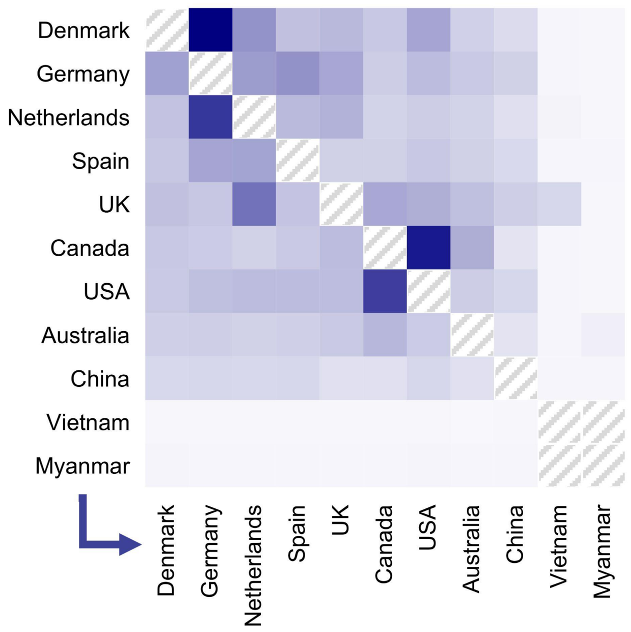


**Figure S18. Variation in rates of between-country transmission of pathogenic lineages.** Heat map showing the relative frequency of transmission between countries across the six most pathogenic lineages based on the analysis shown in Figure 3. These estimates are based on estimated numbers of transmission events for the 6 most common pathogenic lineages using lineage-specific asymmetric models of between-country transmission over time-scaled phylogenies. Cells with diagonal grey lines indicate either identity between countries or country pairs where no lineage is found in both.

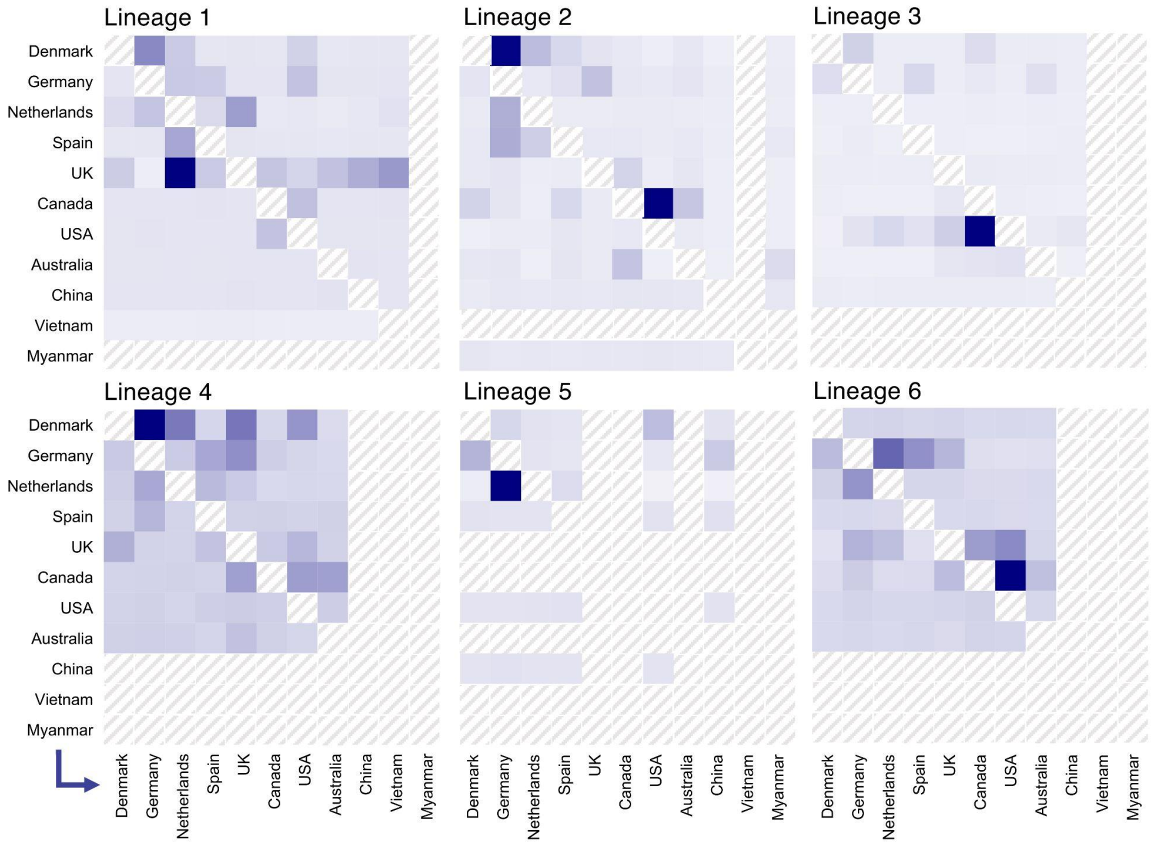


**Figure S19. Variation in rates of between-country transmission for each of the six most common pathogenic lineages.** Heat maps showing the relative frequency of transmission between countries across the six most pathogenic lineages based on the analysis shown in Figure 3. These estimates are based on estimated numbers of transmission events for the 6 most common pathogenic lineages using lineage-specific asymmetric models of between-country transmission over time-scaled phylogenies. Cells with diagonal grey lines indicate either identity between countries or country pairs where the lineage is not found in both countries.

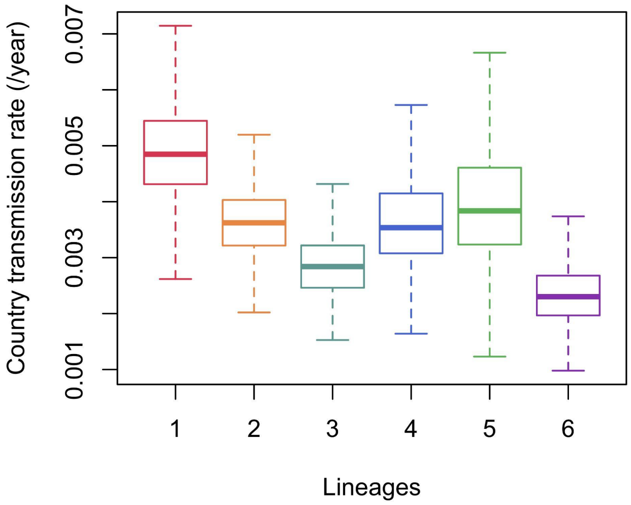


**Figure S20.** Rates of between-country transmission for each of the six most common pathogenic lineages. Rates were estimated using a discrete asymmetric model in BEAST. Interpreting variation in rates of between-country transmission between lineages is made difficult due to variation in the distribution of lineages across countries and variation in the rates of transmission between countries, both of which are likely to have varied over time.

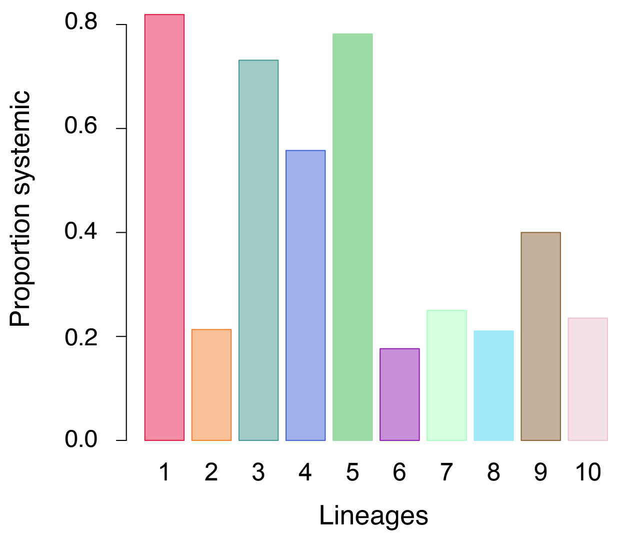


Fisher's exact test results

| | 1 | 2 | 3 | 4 | 5 | 6 | 7 | 8 | 9 | 10 |
|---|---|---|---|---|---|---|---|---|---|---|
| 1 (n=99) | | *** | - | - | - | *** | *** | *** | - | ** |
| 2 (n=95) | | | *** | * | *** | - | - | - | - | - |
| 3 (n=76) | | | | - | - | *** | *** | ** | - | - |
| 4 (n=60) | | | | | - | ** | - | - | - | - |
| 5 (n=56) | | | | | | *** | *** | *** | - | * |
| 6 (n=55) | | | | | | | - | - | - | - |
| 7 (n=24) | | | | | | | | - | - | - |
| 8 (n=30) | | | | | | | | | - | - |
| 9 (n=6) | | | | | | | | | | - |
| 10 (n=17) | | | | | | | | | | |

**Figure S21.** Proportions of disease-associated isolates that are associated with systemic forms of disease in the 10 pathogenic lineages and pairwise comparisons of the frequencies of respiratory and systemic disease isolates using Fisher's exact test with Bonferroni correction for multiple testing. Tests compared the numbers of isolates from each lineage associated with carriage, and respiratory and systemic forms of disease. - indicates p > 0.01, * indicates p < 0.01, ** indicates p < 0.001, *** indicates p < 0.0001. The numbers of isolates with characterised disease-association considered in the analysis are indicated in the row labels.

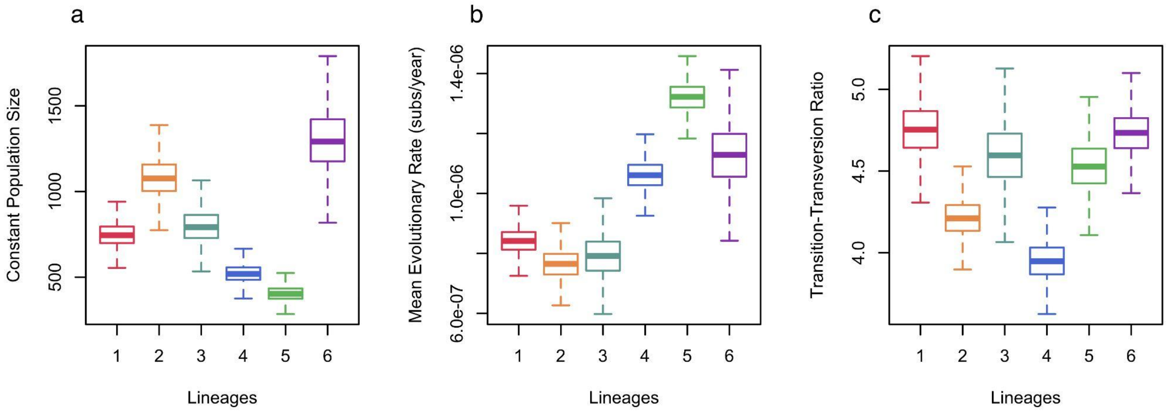


**Figure S22. Estimates of (a) population size, (b) evolutionary rate and (c) transition-transversion rate (kappa) for each of the pathogenic lineages.** This was based on an analysis with BEAST using a strict clock and a constant population size model.

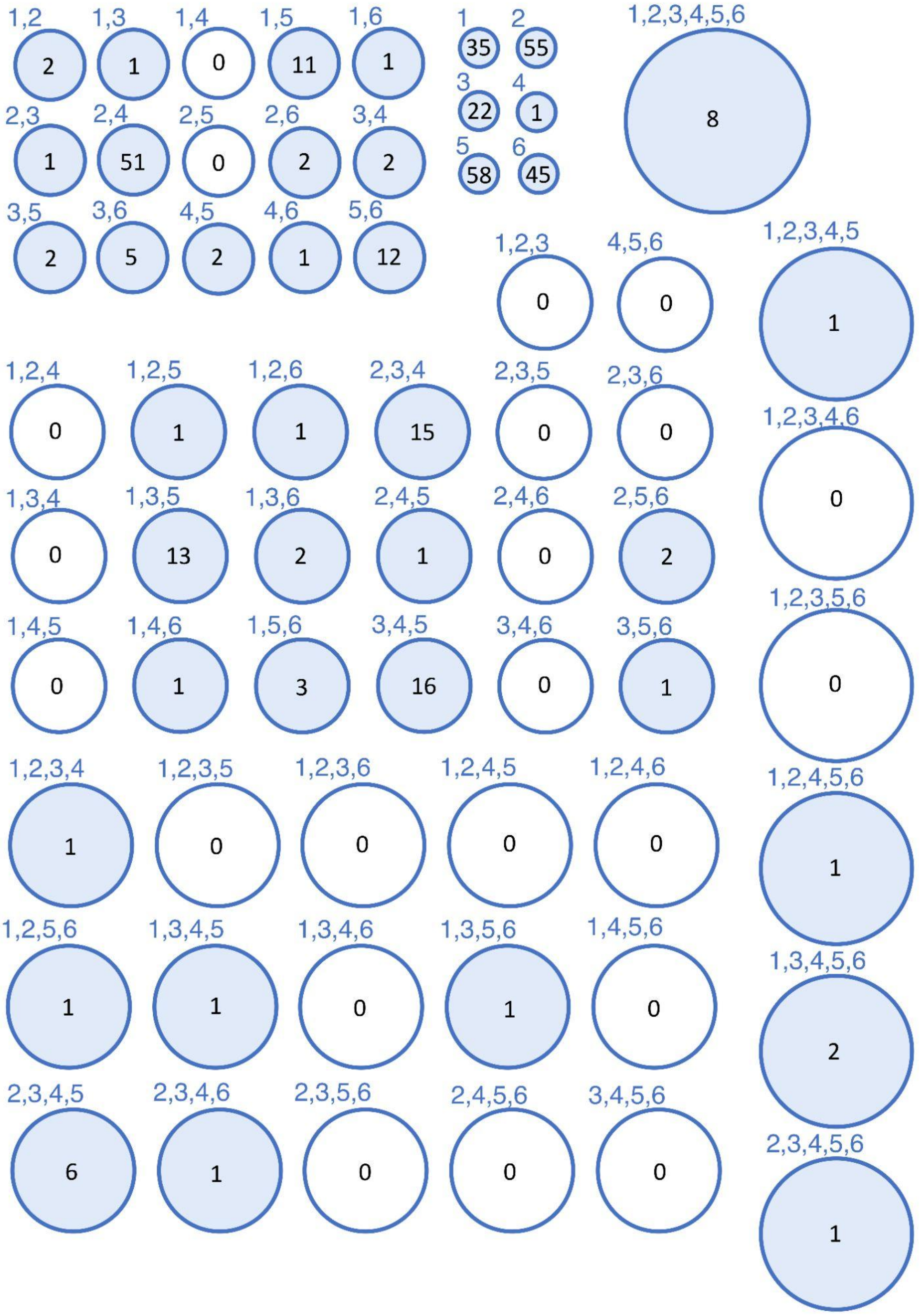


**Figure S23.** The distribution across the 6 most common pathogenic lineages of the 390 genes that are present in >90% of isolates in at least one of the six most common pathogenic lineages and in <10% of isolates outside of the pathogenic clade. Circles represent all possible sets of 6 common pathogenic lineages, with the numbers inside representing the number of genes present in >90% of isolates in each of the lineages in the set and <90% of isolates in each of the six pathogenic lineages outside of the set. Sizes of circles reflect the number of lineages that share these genes.

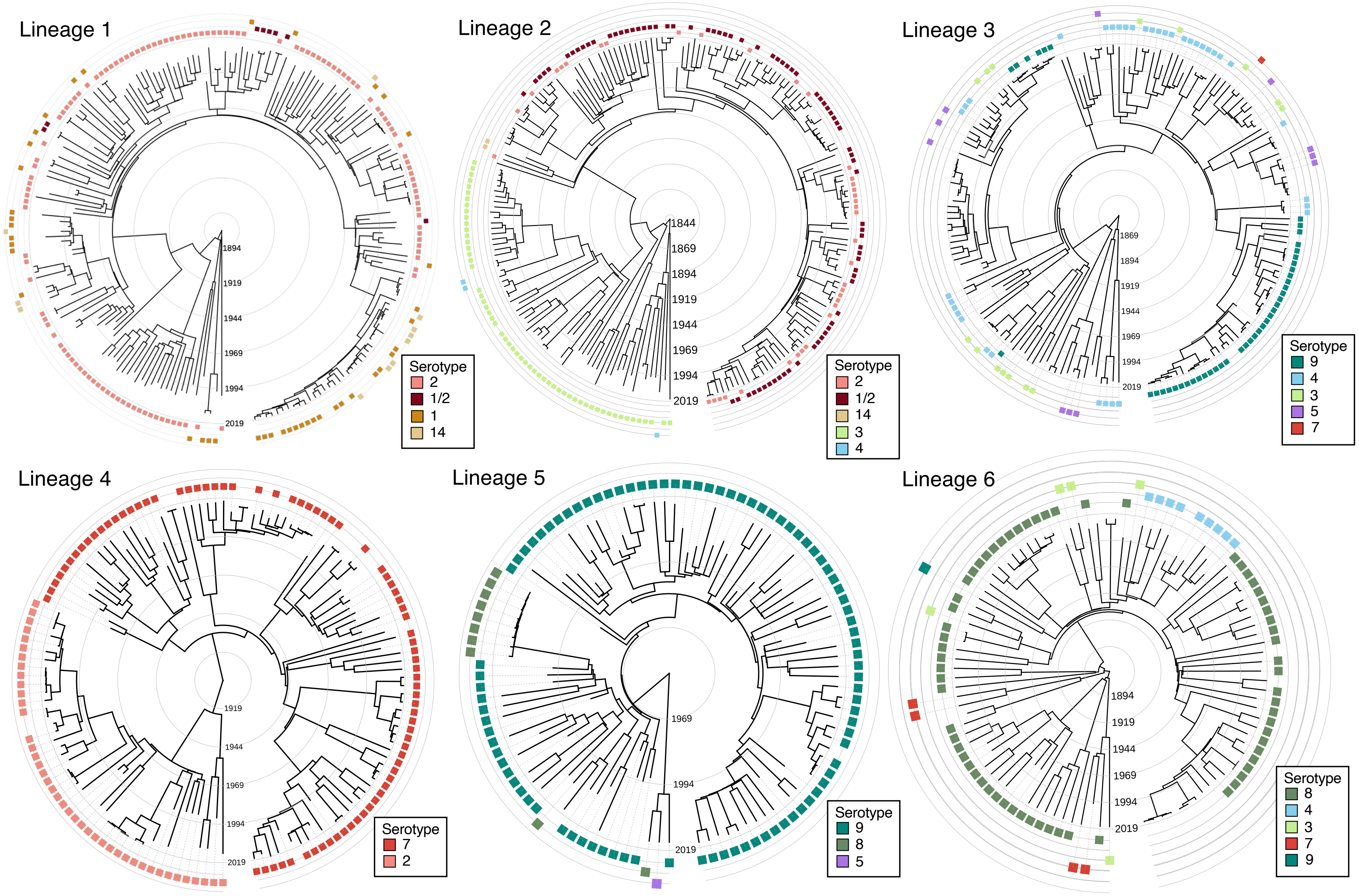


**Figure S24.** Serotypes mapped onto the phylogenies of the six most common pathogenic lineages.

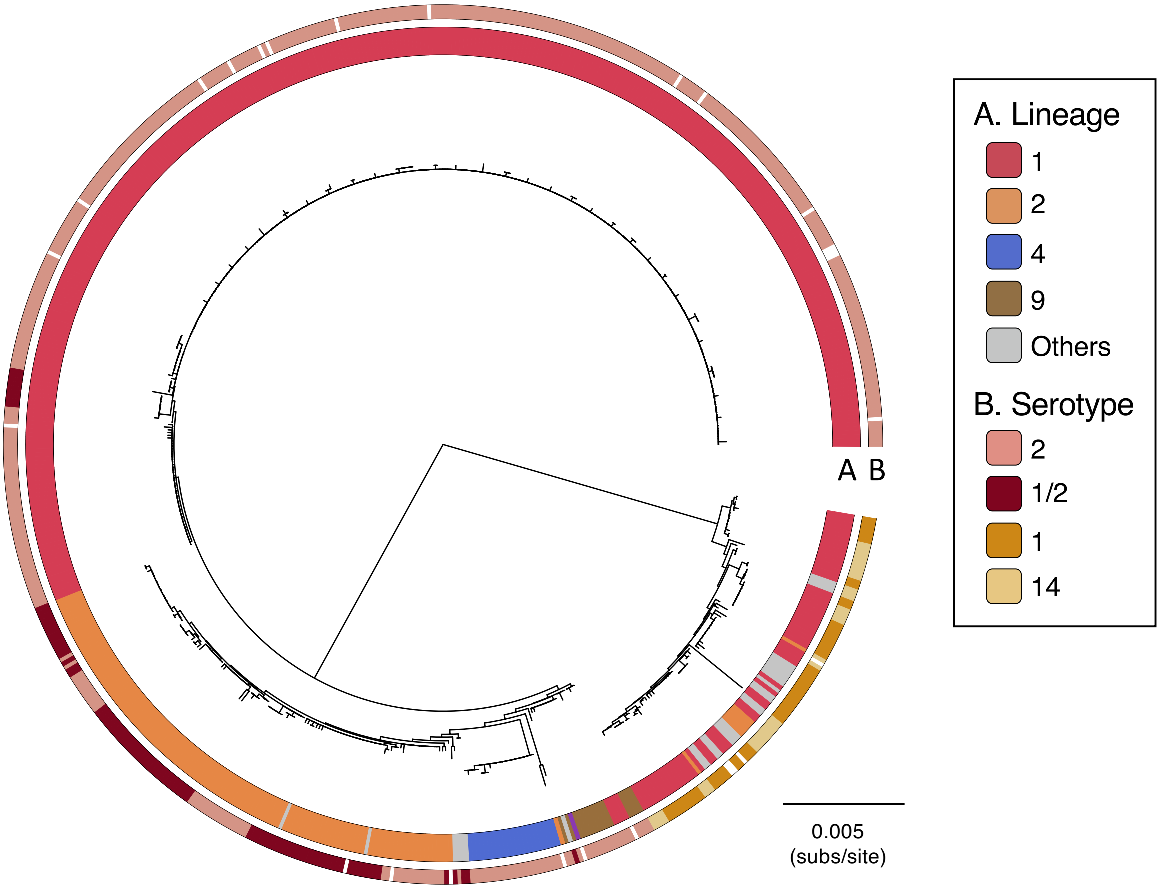


**Figure S25.** A neighbour-joining tree constructed from a concatenated alignment of the genes in the serotype 2 locus (which are shared by the 1/2, 1 and 14 capsular loci). Lineage and serotype are shown in the two outer rings. Isolates with uncertain capsular types are left white. This reveals two divergent clades that each include a pair of serotypes that cannot be distinguished with PCR schemes: 2 and 1/2, and 1 and 14 (Athney *et al.* 2016). The phylogeny suggests that there have been repeated transitions between serotypes 2 and 1/2 and between serotypes 1 and 14.

### Tables

**Table S1.** Isolates in our collection with associated metadata, description of lineage, clade and group, presence/absence of pathogenicity-associated genomic islands and BioProject and assembly IDs.

**Table S2.** Genes associated with pathogenic lineages. These genes were found to be present at >70% higher frequency in pathogenic lineages relative to lineages outside of the pathogenic clade.

**Table S3.** Estimated dates of the origins on the six most common pathogenic lineages.

**Table S4.** Estimated counts and rates of jumps from one country (rows) to another (columns) for each of the six most common pathogenic lineages. Values are median estimates.

**Table S5. Differences in the frequencies of non-clinical isolates from the ten pathogenic lineages across countries in our collection.** To avoid biases due to differences in laboratory protocols in identifying *S. suis* we excluded isolates from outside of the core *S. suis* population. Only isolates sampled from the tonsils or throats of pigs without known *S. suis* disease are included. Due to differences in sampling strategies across our collections from different countries (e.g. geographic and temporal distributions) comparisons across countries should, however, be treated with caution.

**Table S6. Table of genes associated with individual pathogenic lineages.** Genes that are present in >95% of isolates from at least one pathogenic lineage and <5% of isolates from outside of the pathogenic clade.
